# Lysosomal mitochondrial interaction promotes tumor growth in squamous cell carcinoma of the head and neck

**DOI:** 10.1101/2023.06.25.546311

**Authors:** Avani Gopalkrishnan, Nathaniel Wang, Silvia Cruz-Rangel, Abdul Yassin- Kassab, Sruti Shiva, Chareeni Kurukulasuriya, Satdarshan P. Monga, Ralph J DeBerardinis, Kirill Kiselyov, Umamaheswar Duvvuri

**Affiliations:** Department of Otolaryngology, University of Pittsburgh School of Medicine, Pittsburgh, PA; UPMC Hillman Cancer Center, University of Pittsburgh Medical Center, Pittsburgh, PA; Dept of Pharmacology and Chemical Biology, Vascular Medicine Institute, University of Pittsburgh School of Medicine, Pittsburgh, PA; University of Pittsburgh School of Medicine, Pittsburgh, PA; Division of Experimental Pathology, Department of Pathology, Division of Gastroenterology, Hepatology and Nutrition, Department of Medicine, University of Pittsburgh School of Medicine, Pittsburgh, Pennsylvania; Children’s Medical Research Institute and Howard Hughes Medical Institute, University of Texas Southwestern Medical Center, Dallas, Texas; Department of Biological Sciences, University of Pittsburgh, PA; Veterans Affairs Pittsburgh Health System, Pittsburgh, PA

**Keywords:** Lysosomal mitochondria interaction (LMI), TMEM16A, β-catenin, complex I, IACS-10759

## Abstract

Tumor growth and proliferation are regulated by numerous mechanisms. Communication between intracellular organelles has recently been shown to regulate cellular proliferation and fitness. The way lysosomes and mitochondria communicate with each other (lysosomal/mitochondrial interaction) is emerging as a major determinant of tumor proliferation and growth. About 30% of squamous carcinomas (including squamous cell carcinoma of the head and neck, SCCHN) overexpress TMEM16A, a calcium-activated chloride channel, which promotes cellular growth and negatively correlates with patient survival. TMEM16A has recently been shown to drive lysosomal biogenesis, but its impact on mitochondrial function is unclear. Here, we show that (1) patients with high TMEM16A SCCHN display increased mitochondrial content specifically complex I; (2) *In vitro* and *in vivo* models uniquely depend on mitochondrial complex I activity for growth and survival; (3) β-catenin/NRF2 signaling is a critical linchpin that drives mitochondrial biogenesis, and (4) mitochondrial complex I and lysosomal function are codependent for proliferation. Taken together, our data demonstrate that LMI drives tumor proliferation and facilitates a functional interaction between lysosomes and mitochondria. Therefore, inhibition of LMI may serve as a therapeutic strategy for patients with SCCHN.

## Introduction

There are many examples of cancer cells hijacking, enhancing, or otherwise realigning the role normally played by organelles to support increased proliferation ^1, 2^. Alterations in the mitochondrial function can lead to increased energy production, resistance to apoptosis, oxidative stress, and uncontrolled cell growth and division, contributing to the development and progression of cancer ^3^. Dysregulation of lysosomal function can lead to enhanced sequestration and decreased bioavailability of chemotherapeutic drugs, increased degradation of pro-apoptotic proteins leading to suppressed apoptosis, and secretion of the lysosomal enzymes contributing to the degradation of the extracellular matrix enabling cancer cell invasion and metastasis ^4^.

The regulation of bioenergetics in tumor cells has been shown to impact both cellular proliferation and tumor metastasis ^5^. Interestingly, mitochondrial electron transport chain is required for glucose and glutamine metabolism in certain tumors ^6^. Similarly, inhibition of complex I was found to rescue resistance to B-Raf inhibition in melanomas ^7^. These findings suggest that mitochondrial respiration is intimately linked with tumor growth and treatment resistance. However, the exact mechanisms by which mitochondrial respiration is regulated remain unclear.

The synergistic interaction between lysosomes and mitochondria and its role in tumor growth and proliferation has been a topic of recent interest ^8^. Prior investigations have focused on mitophagy as the primary outcome of the interaction between lysosomes and mitochondria ^9^. However, a newer focus has shifted to signaling and functional interaction between these organelles in pathways distinct from mitophagy.

An emerging question is whether regulation of lysosomal biogenesis and mitochondrial function (defined by either biogenesis or respiration) occurs in a coordinated fashion ^10^. Lysosomal biogenesis is regulated by the EB family of transcription factors (including TFEB and MiTF) ^11^. Mitochondrial biogenesis is controlled by a system of transcription factors that are activated by hypoxic signals and other inputs pertaining to redox regulation ^12^. While the mounting evidence suggests interaction between lysosomal and mitochondrial compartments exists ^13, 14^ the impact of this coordination in malignant cells has yet to be consistently explored. The central question pursued in these studies is whether and how the lysosomes and mitochondria functionally interact with each other (biogenesis and function); and whether this interaction can be exploited to target cancer cells.

SCCHN is a devastating disease with overall survival of ∼50% at 5 years. This disease is characterized by amplification of chromosomal band 11q13 in about 30% of cases. These patients have much poorer oncologic outcomes ^15, 16^ . The 11q13 region encodes the TMEM16A calcium-activated chloride channel ^15, 16^ . Overexpression of TMEM16A, partially driven by gene amplification, activates several signaling pathways, including lysosomal biogenesis, that promotes tumor growth ^17^. Therefore, we used this model system to interrogate the interplay between lysosomal biogenesis and mitochondrial bioenergetics.

Using a combination of bioinformatics, human tumor analyses, and *in vivo* murine models, we demonstrate that lysosomal biogenesis is coordinately regulated with the expression of mitochondrial complex I. Mechanistically, we show that lysosomal and mitochondrial biogenesis is co-regulated through ꞵ-catenin/Wnt signaling axis. This coordinated signaling ultimately promotes cellular fitness and tumor proliferation. Pharmacologic inhibition of mitochondrial complex I is particularly effective in tumors that display this coordinate regulation of lysosomal biogenesis and complex I expression. Taken together, these data support the notion that tumor biomarkers can be used to stratify patients who may benefit from tumor-specific complex I inhibition.

## Results

### TMEM16A overexpressing SCCHN exhibit coordinated regulation of lysosomal biogenesis and mitochondrial complex I level and activity

TMEM16A upregulation is a hallmark and a driver of proliferation in many malignancies, including SCCHN ^18^. We previously showed an upregulation of lysosomal flux comprising both lysosomal biogenesis and exocytosis in human cancers, particularly in the context of TMEM16A upregulation^17^. Therefore, we sought to determine whether TMEM16A drives a synergistic relationship between lysosomes and mitochondria in malignant cells.

We use a combination of evolutionary rate covariation (ERC) and gene expression analysis using a publicly available cancer cell line database of gene expression for an unbiased prediction of the functional content of lysosome-mitochondria interaction in TMEM16A-upregulating SCCHN models. ERC is a computational approach that presumes shared evolutionary pressure and, therefore, a collective evolutionary history of proteins that are involved in functional interactions ^19^. Such histories can be tracked and correlated by calculating the divergence of amino acid or gene sequence pairs in several organisms in a clade or a set of clades. Proteins whose evolutionary histories correlate are candidates for functional interaction, and these correlations do not rely on prior empiric data.

We used a publicly available ERC portal https://csb.pitt.edu/erc_analysis/ to study the coevolution between known mitochondrial and lysosomal proteins in mammals. The analysis was based on the Mitocarta 3 database (1136 proteins) https://www.broadinstitute.org/mitocarta/mitocarta30-inventory-mammalian-mitochondrial-proteins-and-pathways and a published lysosomal proteome (417 proteins)^20^. The mitochondrial proteins were divided into groups based on their location in the organelle (outer membrane and intermembrane). Similarly, we segregated the lysosomal proteins into those containing (TMD+) and not containing transmembrane domains (TMD-). The coevolution between proteins was calculated as the correlation between their percent identity values in 49 species of mammals (Supplementary Table ST1 and ST2). Table 1 lists the protein types and specific coevolving proteins using coevolution with 10% of the corresponding organelle’s proteome as a threshold.

**Table 1:**
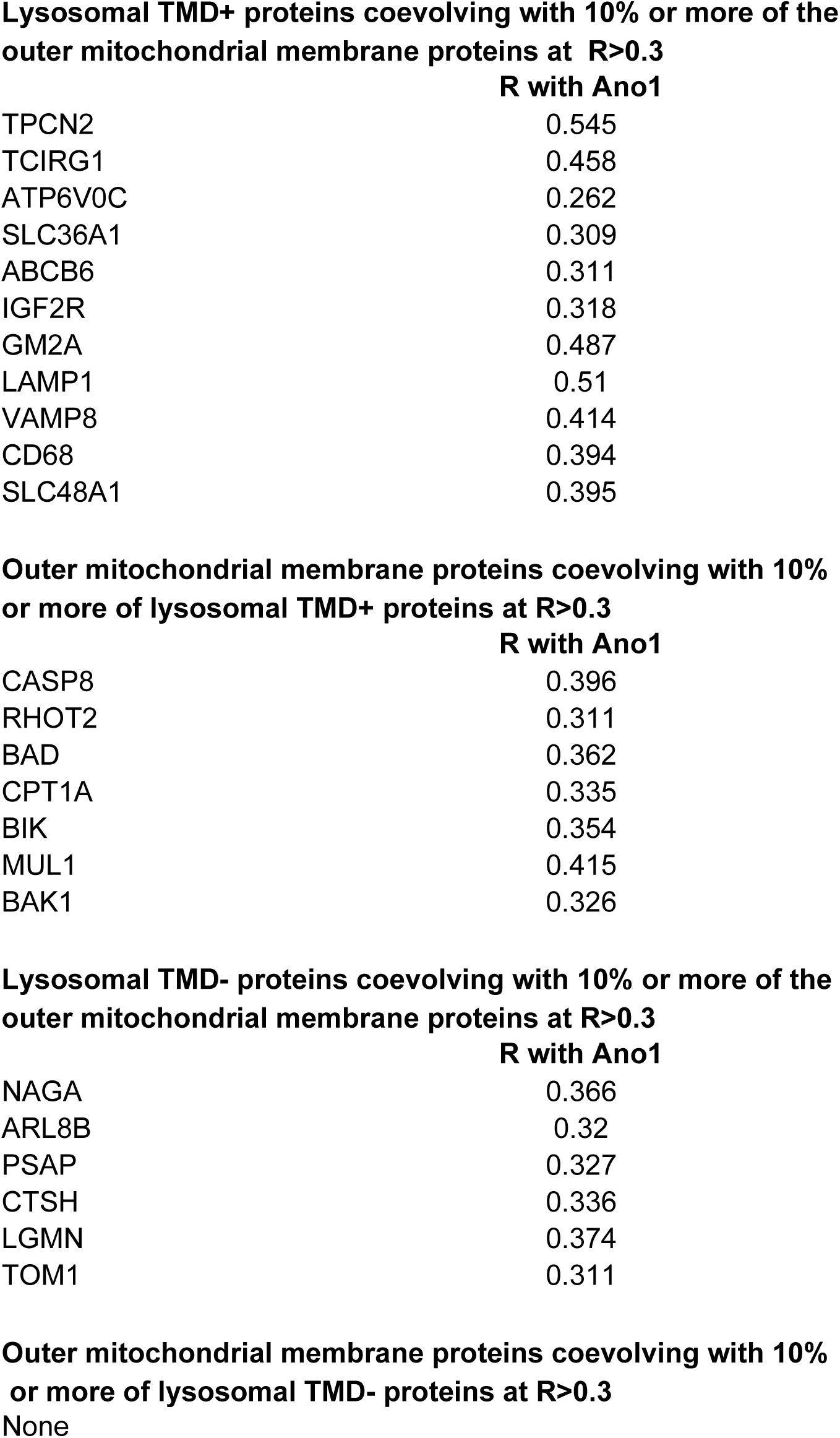
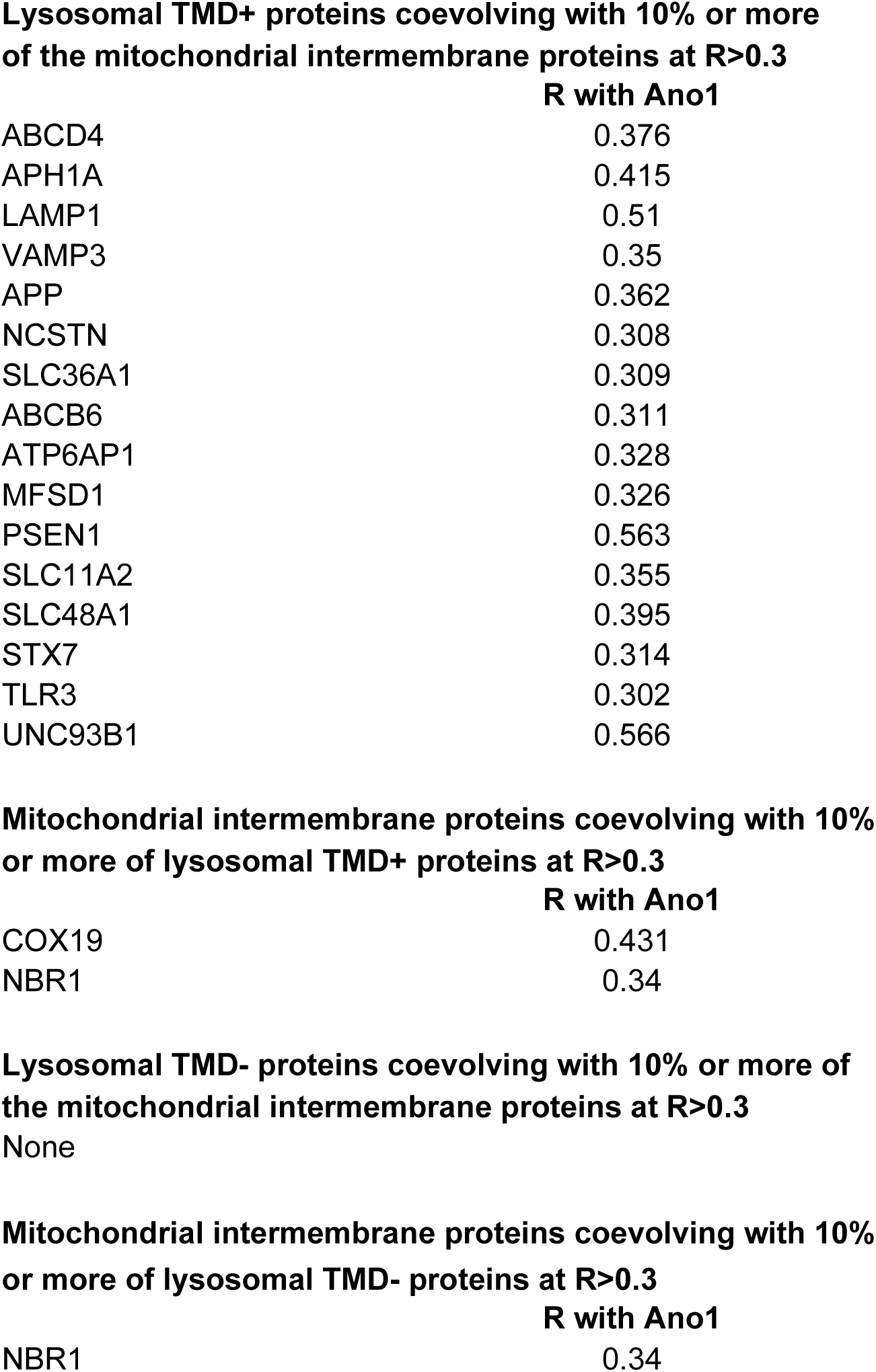
Lysosomal and mitochondrial components that show significant coevolution (at ERC R≥0.3) with 10% of the corresponding organellar proteome and R≥0.3 correlation with TMEM16A expression in upper aerodigestive cancer cell lines.

A detailed analysis of these protein classes identifies several potential functional complexes shared by the outer mitochondrial membrane and lysosomal proteins, including proteins involved in ion exchange, mitochondrial positioning, and fusion/fission. This coevolution indicates a close functional (but not necessarily physical) engagement between lysosomes and mitochondria. Next, we sought to understand if this interaction is enriched in tumors that are putatively more aggressive. We used TMEM16A expression as a surrogate of tumor aggressiveness ^21^. Therefore, we parsed Cancer Dependency Map database, Expression 22Q2 dataset, focusing on cancer cell lines of upper aerodigestive origins (58 lines). This approach yields a set of genes that co-evolve and whose expression is co-regulated in the context of TMEM16A expression (Fig. 1A). Interestingly, these analyses revealed a specific co-regulatory effect on genes coding for the mitochondrial complex I components.

**Figure 1.**
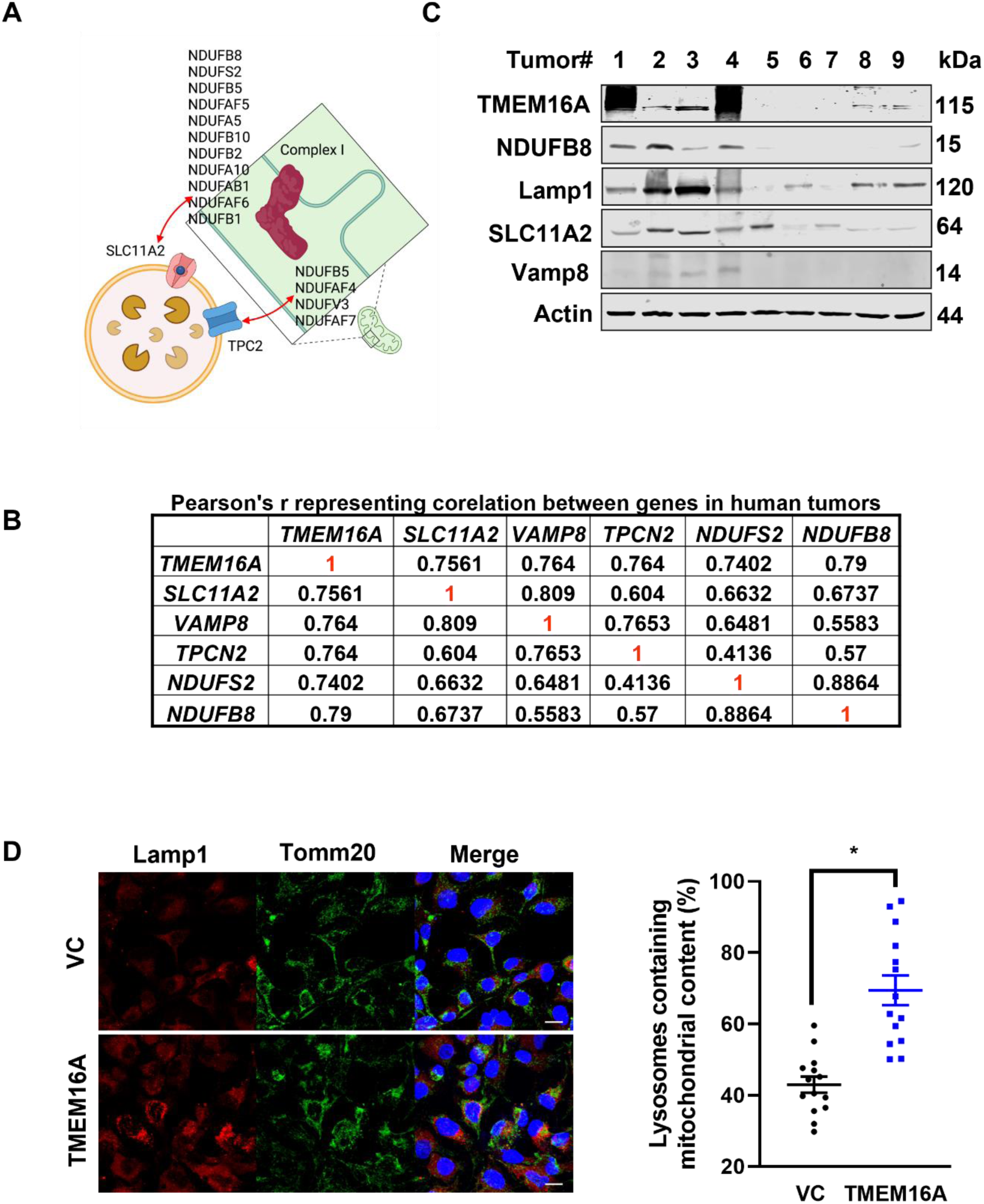
TMEM16A overexpressing SCCHN exhibit coordinated regulation of lysosomal biogenesis and mitochondrial complex I. ERC analysis reveals that lysosomal proteins and mitochondrial complex I proteins co-evolve and are co-expressed in squamous cell carcinoma of the head and neck (A). Immunoblots of human tumor tissue show that these proteins are co-expressed in tumors that display overexpression of TMEM16A (B). Gene expression is also co-regulated in a larger cohort of human SCCHN tumors (N=24). Correlation coefficients of regression analyses of various pairs of genes are shown in. These correlations all demonstrated significance (p < 0.05) (C). Immunofluorescence analyses demonstrates co-localization of lysosomes with mitochondria in the context of TMEM16A overexpression (D).

We next sought to confirm these findings using a cohort of human SCCHN tumor tissues. We analyzed the expression of various lysosomal and mitochondrial genes using qPCR, followed by a correlation analysis. The correlation coefficients from these analyses are shown in Fig. 1B and Supplementary Fig. S1A, B. Interestingly, we found a significant correlation between the expression of genes coding for lysosomal and mitochondrial complex I proteins, in the context of TMEM16A overexpression (Fig. 1C). We observed a particularly robust induction of complex I genes upon TMEM16A expression (Supplementary Fig. S1C). However, there was a lack of correlation between TMEM16A, and genes related to complex III and IV; suggesting that the link between lysosomal biogenesis and mitochondrial function is specifically mediated by complex I (Supplementary Fig. S1D-G).

We have previously shown that TMEM16A-induced lysosomal biogenesis causes a collateral increase in lysosomal exocytosis ^17^. Therefore, we sought to determine if other lysosomal functions like autophagy are also increased. Using EM, we identified increased autophagic vacuoles (defined as double membraned intracellular vacuoles) in TMEM16A overexpressing tumors (Supplementary Fig. S1D, E). We further expanded our investigations to determine whether the observed co-expression translated to a potential physical interaction between lysosomes and mitochondria. Expectedly, lysosomal biogenesis (triggered by TMEM16A overexpression) leads to increased co-localization between these organelles (Fig. 1D). Mitophagy is a process by which effete mitochondria are degraded.

Given the observed co-localization between lysosomes and mitochondria, we determined the effect of mitophagy in our cells. Upon treatment with the electron transport chain uncoupler FCCP, we noted an induction of parkin expression (indicative of mitophagy). However, there was no increase in parkin induction in the context of TMEM16A overexpression. This suggests that TMEM16A does not directly impact mitophagy (Supplementary Fig. S1F), suggesting that the observed co-localization is not due to increased mitophagy. Taken together, these data demonstrate that lysosomal biogenesis is coordinately regulated with the expression of mitochondrial complex I proteins, independently of mitophagy.

### TMEM16A promotes mitochondrial biogenesis

Our prior studies have shown that TMEM16A promotes lysosomal biogenesis in SCCHN models ^17^. Here, we sought to determine if TMEM16A promotes mitochondrial biogenesis. Mitotracker staining in cells engineered to overexpress TMEM16A evidenced increased mitochondrial mass (Fig. 2A). We find that several transcription factors that have a role in mitochondrial biogenesis, redox balance, and defense against oxidative damage, including *PGC1α* and *NRF2* are amplified in TMEM16A-overexpressing OSC19 (Fig. 2B). This is in concert with the prior evidence of higher baseline levels of reactive oxygen species (ROS) in TMEM16A-overexpressing SCCHN models ^17, 22^. TMEM16A-promoted mitochondrial biogenesis is not only associated with increased expression of genes linked with mitochondrial antioxidants (*SOD1*, *SOD2* and *GPX*) and regulators of mitochondrial respiration (cytochrome oxidase and citric synthase) (Fig. 2B).

**Figure 2:**
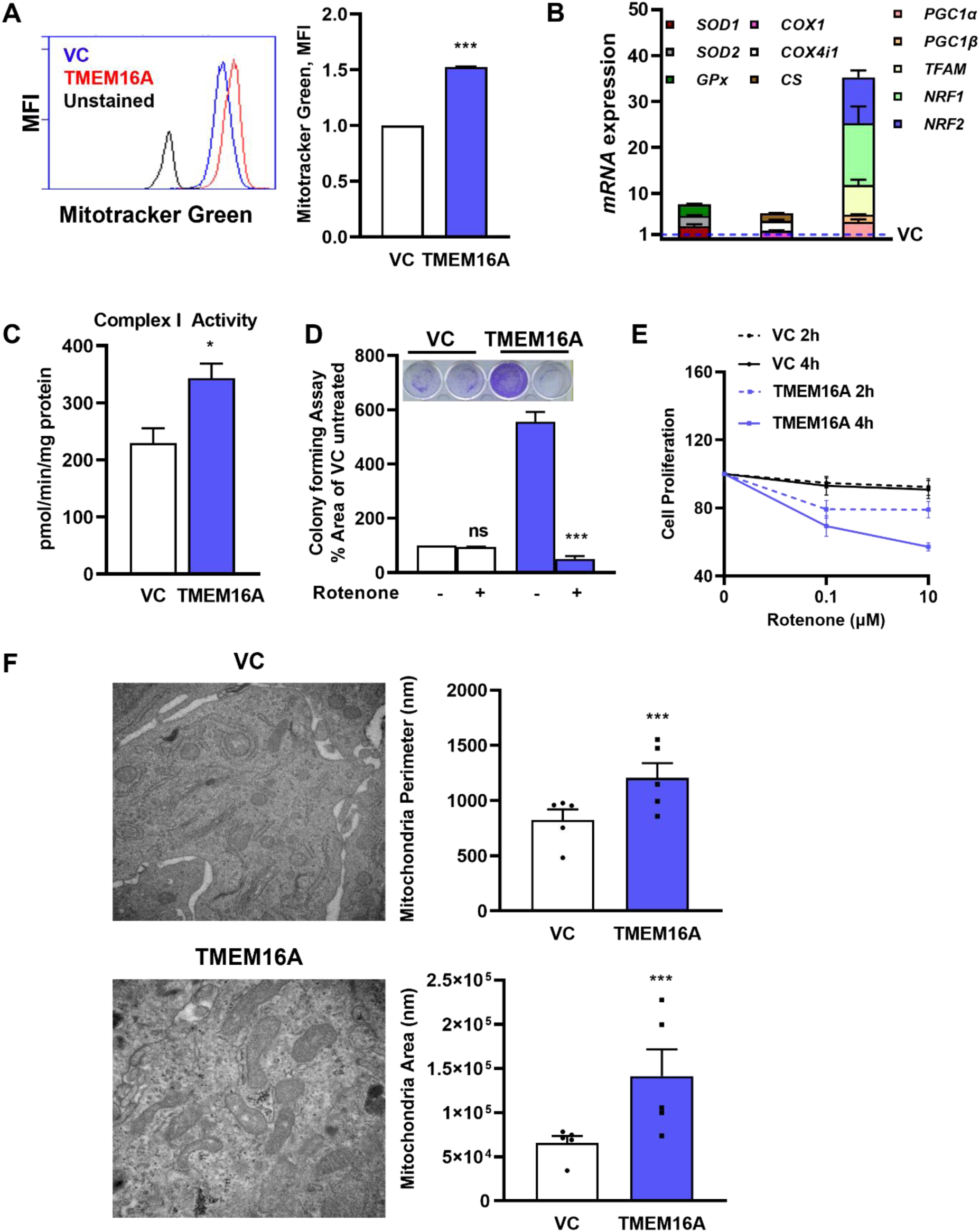
TMEM16A promotes upregulation of mitochondrial mass and biogenesis. OSC19 cells were engineered to stably overexpress TMEM16A. These cells were subjected to flow cytometry with Mitotracker dye to measure mitochondrial mass. TMEM16A overexpression increased Mitotracker staining (A). Gene expression analyses revealed a profound overexpression of genes associated with mitochondrial biogenesis in these cells (B). Consistent with prior data, the activity of mitochondrial complex I was increased upon TMEM16A overexpression (C). Vector control (VC) or TMEM16A overexpressing cells were treated with diluent or rotenone for 2-4h and assessed for cell viability using WST assays (D). Similarly, cells were treated with diluent or rotenone and assessed for cell viability using colony formation assays (E). Tumor xenografts were generated by inoculating nude mice with the respective cells. Tumors were harvested and subjected to EM analyses to measure mitochondrial perimeter and area (F).

Proteomic and biochemical analyses reveal that the expression of complex I proteins is increased upon TMEM16A expression, and this is correlated with an increase in complex I activity (Fig. 2D-F). We extended these findings to determine if knock-down of TMEM16A abrogated the phenotypes described above. Indeed, genetic knock-down of TMEM16A in SCCHN cells that express endogenously high levels of the gene (HN30), led to significant decrease in mitochondrial biogenesis, mass and complex I activity (Supplemental Fig. S6A-D).The upregulation of mitochondrial biogenesis in TMEM16A-overexpressing OSC19 cells is reflected in the pronounced dependence of these cells’ proliferation on mitochondrial function. Suppressing the mitochondrial function using rotenone (Fig. 2D, E) and complex I inhibitor IACS-10759 (Fig. 5A), significantly decreased proliferation of specifically TMEM16A-overexpressing cell line. As an additional control, IACS-10759 inhibited the enhanced ATP production and mitochondrial membrane potential in the TMEM16A-overexpressing OSC19 cell line (Fig. 5B, C). Similarly, using xenograft models, we find that TMEM16A overexpression leads to increased mitochondrial mass, as measured by mitochondrial area and perimeter on electron microscopic images (Fig. 2F).

### Mitochondrial biogenesis is regulated by ꞵ-catenin signaling in the context of TMEM16A overexpression

We have previously shown that upregulated lysosomal biogenesis in TMEM16A-overexpressing SCCHN models is driven, at least in part, by ꞵ-catenin signaling ^17^. We and other have shown that ꞵ-catenin can promote both lysosomal biogenesis ^17, 23^, and mitochondrial biogenesis ^24^ in certain cell types. However, these effects have not been as well studied in squamous cell carcinoma. We posit that TMEM16A drives a signaling network that promotes lysosomal biogenesis and thereby facilitates enhanced lysosome-mitochondrial signaling. Therefore, we sought to interrogate the mechanism(s) that regulate this interaction.

We have previously shown that lysosomal biogenesis is driven by ꞵ-catenin signaling^17^. We postulated that ꞵ-catenin signaling serves as a ‘linchpin’ to regulate both processes. We therefore measured ꞵ-catenin activity in cells that were engineered to overexpress TMEM16A. As expected, we found increased ꞵ-catenin activity in the context of TMEM16A overexpression (Fig. 3A, Supplemental Fig. 6E). ꞵ-catenin inhibitor PRI-724 induced a significant decrease in ꞵ-catenin activity in TMEM16A overexpressing cells, but not in control cells (Fig. 3A). From a mechanistic perspective, we noted that treatment with PRI-724 significantly reduced the expression of transcription factors that drive mitochondrial biogenesis (Fig. 3B). This was accompanied by a decrease in ATP production (Fig. 3C), consistent with the hypothesis that ꞵ-catenin activity drives mitochondrial function. TMEM16A cells were also noted to be sensitive to cell proliferation via PRI-724 (Fig. 3D).

**Figure 3:**
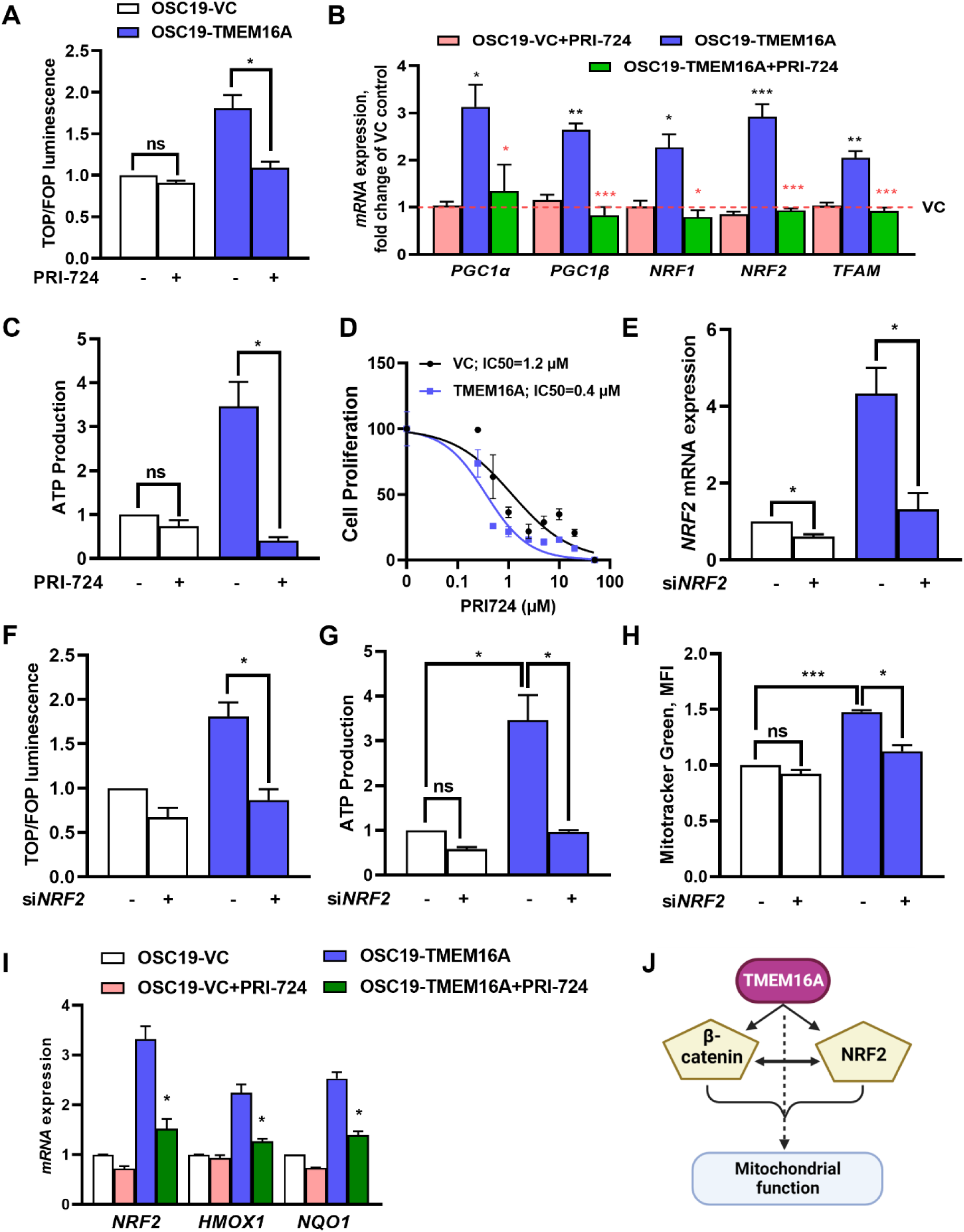
βcat/NRF2 coactivation in TMEM16A is pivotal in regulating mitochondrial function. TMEM16A overexpression promotes ꞵ-catenin activity, that is abrogated by treatment with ꞵ-catenin inhibitor PRI-724 (2µM, 24h) (A). The effect of PRI-724 treatment on markers of mitochondrial biogenesis is shown in (B). The effect of PRI-724 treatment on ATP production was measured using the ATPlite assay (C). The effect of PRI-724 treatment on cell proliferation, as determined by WST assay is shown in (D). Transient knockdown of NRF2 is measured by qPCR (E). NRF2 knockdown depletes βcatenin activity and mitochondrial function (F, G) and mitochondrial mass (H). βcatenin inhibition via PRI-724 targets *NRF2* signaling pathway (I). The putative model for TMEM16A induced mitochondrial biogenesis (J).

Since, ꞵ-catenin mutations are not frequently encountered in squamous cell carcinoma, enhanced ꞵ-catenin activity should be driven by ligand dependent signaling. Therefore, we used an inhibitor of Porcupine (LGK-974, which abrogates the release of WNT ligands) as an indirect inhibitor of ꞵ-catenin. As expected, treatment with LGK-794 induced a significant decrease in ꞵ-catenin activity, ATP production and genes of mitochondrial as well as lysosomal biogenesis Supplemental Fig. S2A-E). However, cell viability was not significantly impacted by treatment with LGK-794, suggesting that other signaling pathways downstream of WNT/ꞵ-catenin signaling may be impacting tumor cell survival (Supplemental Fig. S2F & S6G).

NRF2 pathway and ꞵ-catenin coactivation have been implicated in hepatocellular cancer^25^. Since our data indicates increased activity of ꞵ-catenin and NRF2 transcript, we sort to understand if these pathways cooperate to enhance TMEM16A induced mitochondrial biogenesis. Transient knockdown of NRF2 depleted ꞵ-catenin activity and reduced mitochondrial function and mass (Fig. 3E-H). Interestingly, PRI-724 detracted the transcript expression of NRF2 signaling genes (Fig. 3I). These data demonstrate cooperation between the β-catenin signaling and NRF2 pathway in regulating mitochondrial function (Fig. 3J).

### Interaction between lysosomes and mitochondrial complex I regulate cancer cell viability

Based on the data described above, we postulated that a coordinated signaling between lysosomes and mitochondria regulates cellular biogenetics (Fig. 4A). We first tested the hypothesis that inhibition of mitochondrial complex I disrupts lysosomal biogenesis. Treatment of SCCHN with the complex I inhibitor IACS-10759 led to a significant decrease in lysosomal biogenesis, as assessed by the expression of integral lysosomal genes (Fig. 4B). Interestingly, treatment with inhibitors of complex IV (ADDA5, a cytochrome oxidase inhibitor), complex III (Atovoquone) or oxidative phosphorylation uncoupler FCCP did not significantly impact lysosomal biogenesis (Supplemental Fig. S4). These data along with our bioinformatic analysis strongly suggest the interaction between lysosomes and mitochondria is driven by complex I. A corollary of this observation is that inhibition of the integral lysosomal genes involved in lysosome mitochondria axis in TMEM16A overexpressing SCCHN, as determined from the ERC analysis should impact mitochondrial oxidative phosphorylation. Transient knockdown of lysosomal genes *TPCN2, SLC11A2* and *VAMP8* reduced ATP production (Supplemental Fig. 5A, B, Fig. 4C, D) and impacted the cell growth as observed in colony forming assay (Fig. 4E). Additional inhibition of complex I via IACS-10759 together with knockdown of lysosomal genes did not further deplete ATP, suggesting that that the said mechanism is specific to complex I (Supplemental Fig. 5C-E).

**Figure 4:**
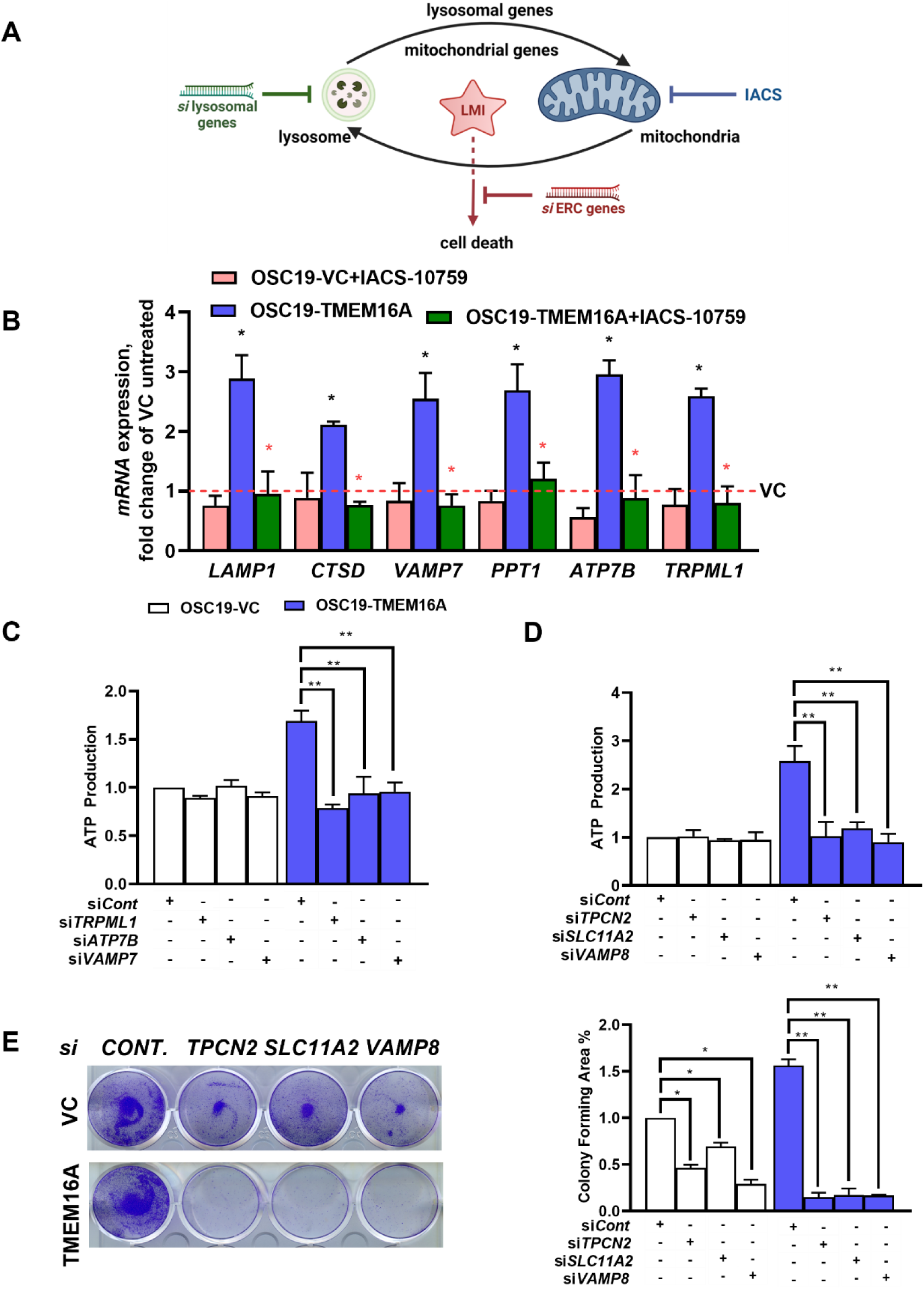
Interaction between lysosomes and mitochondrial complex I regulate cancer cell viability. A proposed model for the interaction is shown in (A). Treatment with the complex I inhibitor, IACS-10759 reduces lysosomal biogenesis (B). Knock-down of lysosomal and ERC genes leads to a reduction in ATP production (C, D) and cell proliferation, as measured by colony formation (E).

The upregulation of mitochondrial biogenesis persisted in the natively high-expressing HN30 cell line (human squamous cell carcinoma of the oral cavity) (Supplementary Fig. 7D). Supplementary Fig. 7G and H show additional evidence for IACS-10759 effects. This drug is in current clinical trials for several cancer types ^26,27^although its effects in the SCCHN context have not been established, which motivated the next set of experiments.

### TMEM16A-driven tumor xenograft models are sensitive to complex I inhibition

To generate a mouse *in vivo* SCCHN model, we subcutaneously injected C57 BL/6J mice with control (VC; Vector Control) and TMEM16A-overexpressing syngeneic MOC1 (squamous cell carcinoma of the mouse oral cavity) cell lines and documented the development of tumors. Mice were treated with 10mg/kg body weight IACS-10759 for 8 consecutive days. The tumor volume data in Fig. 5D and Supplemental Fig. S6A, B show enhanced suppression by IACS-10759, additionally evidenced by decreased Ki67 staining in the TMEM16A IACS-10759 treated tumors (Supplemental Fig. 6C, D). Fig. 5E-G shows the suppressive effects of IACS-10759 using high-TMEM16A expressing patient-derived xenografts (PDX) implanted into immunocompetent NOD SCID mice. Clinical characteristics of the patient (to form PDX) have been tabulated in Supplementary Fig. 6F. Taken together, these data show the unique dependence of TMEM16A-overexpressing SCCHN models on mitochondrial function.

**Figure 5:**
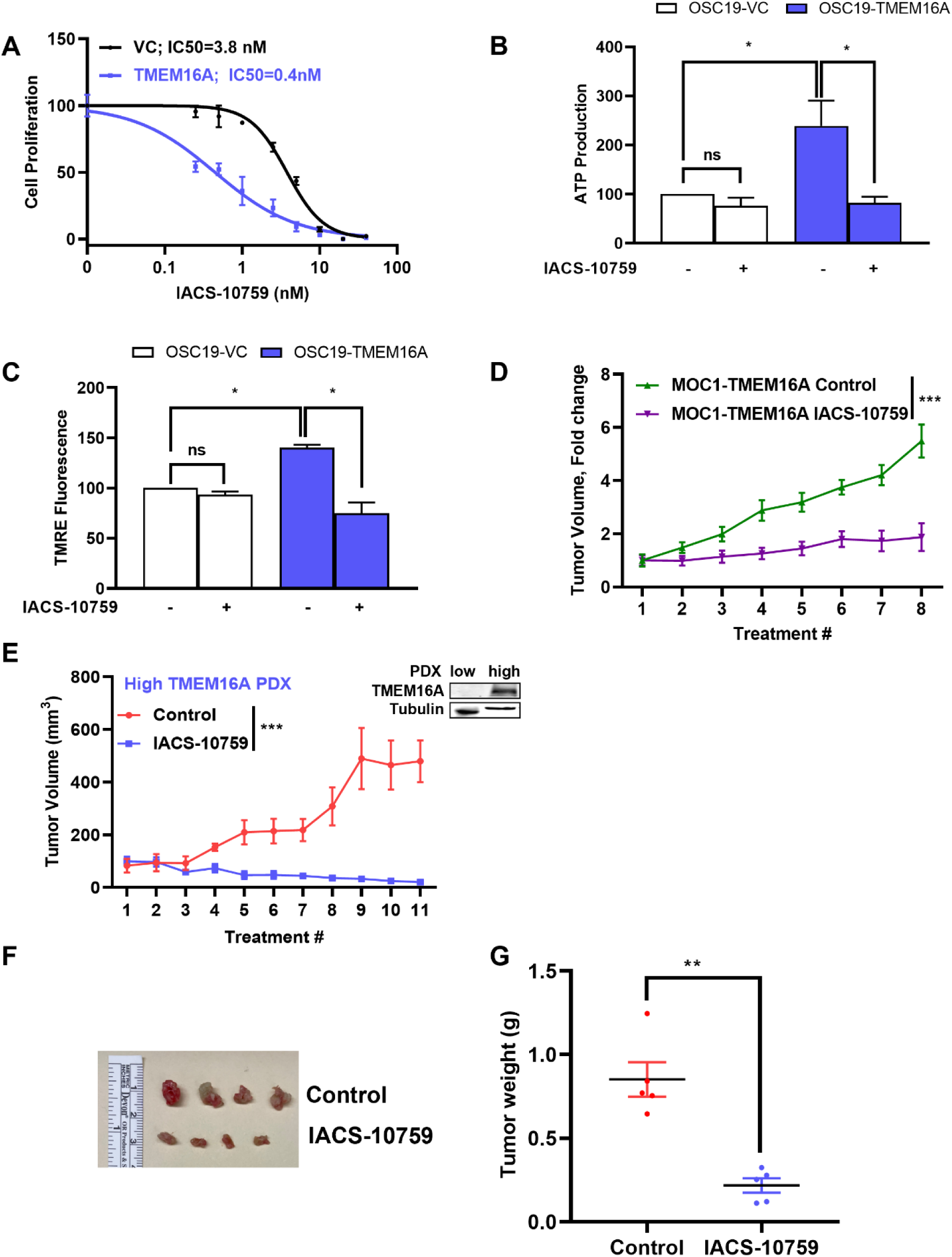
Complex I inhibition sensitizes TMEM16A-driven tumor xenograft models. Control and TMEM16A overexpressing cells were evaluated for sensitivity to the complex I inhibitor, IACS-10759 (A). The effect of IACS-10759 on ATP production and TMRE fluorescence is shown in (B, C). The effect of IACS-10759 on subcutaneous tumors generated from the syngeneic murine SCCHN cell line (MOC1) is shown in (D). A patient-derived xenograft (PDX) was treated with IACS-10759 to demonstrate the anti-tumor activity (E). Representative tumor pictures and tumor weights after necropsy are shown in (F, G).

Since TMEM16A drives cancer cell proliferation ^21^, we wanted to test if these changes could be replicated in conditions of increased cellular proliferation that are independent of TMEM16A ^28, 29^. To test this, we used a transiently overexpressed mutant version of PI3KCA (PI3KCA-H1047R), an oncogenic mutation that is frequently encountered in SCCHN. Interestingly, overexpression of PI3KCA-H1047R did not consistently induce mitochondrial biogenesis, although we noted a significant increase in the expression of the transcription factor TFAM (Supplementary Fig 8A, B). Similarly, we did not observe any increase in lysosomal biogenesis (Supplementary Fig 8C). We confirmed these data by overexpressing EGFR, a known SCCHN oncogene. EGFR overexpression did not promote mitochondrial or lysosomal biogenesis (Supplementary Fig 8D-F). Taken together, these data suggest that LMI is specifically driven by TMEM16A overexpression.

## Discussion

Many of the mechanism(s) that regulate tumor cell growth and proliferation remain elusive despite extensive investigations ^30^. In addition to canonical signaling pathways that regulate cellular proliferation, intracellular organelles also impact cell growth and proliferation ^31^. Indeed, several important signaling molecules like mTORC1, AMPK, GSK3 localize to the lysosomal membrane ^32^. We and others have studied the role of lysosomes in the regulation of cell proliferation since downstream signaling from the lysosomal membrane promotes tumor cell proliferation ^1, 17, 33, 34^.

Lysosomal biogenesis is regulated by the CLEAR network of genes. However, the upstream signaling nodes that regulate the CLEAR network may also have a redundant role in regulating cell proliferation. An important question in the field is how do lysosomes synergize with other organelles in the context of tumor proliferation?

Our data provide evidence that lysosomal-mitochondrial interaction mediates cell growth and proliferation in SCCHN. This interaction is facilitated by the expression of the proto-oncogene TMEM16A. Upon overexpression of TMEM16A, lysosomal and mitochondrial biogenesis are upregulated in a coordinated fashion leading to improved cellular fitness. Specifically, genes that encode components of mitochondrial complex I are enriched in our co-expression analyses. This suggests that there may be a unique survival advantage upon overexpression of mitochondrial complex I proteins. Interestingly, TMEM16A expression seems to promote dependence on mitochondrial respiration in our model. However, lysosomal biogenesis appears to serve a more general role.

Mechanistically, TMEM16A overexpression drives β-catenin signaling, which, in turn, regulates both lysosomal and mitochondrial biogenesis. The effect of β-catenin activation on lysosomal and mitochondrial biogenesis has not been previously reported in SCCHN. SCCHN, unlike other malignancies, does not harbor a high level of β-catenin mutations ^35^. Therefore, β-catenin activation is likely driven by in ligand-dependent manner. Thus, we used an inhibitor of WNT ligand secretion (LGK-974) as a surrogate for β-catenin inhibition. While LGK-974 did inhibit β-catenin activation, it did not induce preferential cell death in our models. However, prior data indicates that treatment with a direct β-catenin inhibitor (PRI-724) demonstrated differential cell death in the context of TMEM16A overexpression ^17^. This suggests that WNT ligand-independent pathways may activate β-catenin signaling and promote cell growth. Nonetheless, β-catenin inhibition induces cell death while ameliorating mitochondrial and lysosomal biogenesis. Furthermore, β-catenin and NRF2 coordinate their activation which drives mitochondrial function, mechanism that has not been shown in the context of SCCHN. In the context of hepatocellular carcinomas, this cooperation has been reported to be relevant to redox balance and tumor development^36^.

Taken together, our data implicate TMEM16A-induced β-catenin activation as an anchor that regulates lysosomal and mitochondrial biogenesis in a coordinated fashion. This interaction between lysosomes and mitochondria promotes tumor cell proliferation. Interestingly, knock-down of specific lysosomal genes reduces ATP production and cell viability in a complex I dependent manner. Knock-down of TPCN2 had the greatest effect on ATP production and cell proliferation (Fig. 4). Furthermore, treatment with a complex I inhibitor did not induce an additional reduction of ATP, suggesting that the LMI is specific to mitochondrial complex I. In concert with this postulate, complex III nor complex V inhibition did not impact lysosomal biogenesis. To the best of our knowledge, this is the first mechanistic description of how lysosomes affect mitochondrial respiration.

It has not escaped our attention that these phenomena may reflect the cellular changes driven by non-specific cellular proliferation. However, expression of constitutively active PI3KCA mutant (PI3KCA-H1047R) or overexpression of EGFR did not promote lysosomal biogenesis or LMI. This provides further evidence to suggest that the TMEM16A/b-catenin axis specifically promotes LMI. However, we cannot completely exclude the possibility that other signaling pathways that promote tumor cell proliferation may also enhance LMI. Further work is needed to fully elucidate the molecular networks that promote/ regulate LMI.

Our data highlight the importance of lysosomal-mitochondrial interaction and how it promotes tumor growth. This is a novel description that highlights how intra-cellular communication between organelles is promoted through intracellular signaling pathways. From a translational perspective, our dissection of the TMEM16A/β-catenin-LMI phenotype implicates β-catenin as a potential molecular target in carcinomas that overexpress TMEM16A, independently of β-catenin mutation status. The anti-tumor activity of PRI-724 in our pre-clinical models provides a strong rationale upon which clinical trials can be initiated to improve treatment for patients with malignancies that are driven by β-catenin signaling independently of mutation status. This could create a new paradigm to treat these patients.

## Materials and Methods

### Cell lines

All cell lines were STR authenticated and confirmed mycoplasma-negative using the e-Myco™ Mycoplasma PCR Detection Kit (Boca Scientific, Inc.). Human tongue squamous cell carcinoma OSC19 were grown in high glucose DMEM supplemented with 10% FBS. OSC19 were transduced with retroviral pBABE-puromycin control or TMEM16A plasmid as previously described here ^37^. Mouse oral squamous cell carcinoma MOC1 cells were grown in Iscove’s Modified Dulbecco’s Medium (IMDM)/Ham’s F-12 Nutrient Mixture 2:1, 5% FBS, 1% penicillin/streptomycin mixture, 5 mg/mL insulin, 40 mg hydrocortisone, 5 mg epidermal growth factor (EGF). MOC1 cells were engineered to overexpress TMEM16A or negative using lentivirus system. After viral infection, cells were selected in puromycin for atleast 2 weeks. TMEM16A expression in the stable cells was confirmed by qPCR.

### siRNA Transfection

About 70% confluent cells in a 6-well plate, were transfected with either 10 µM control (QIAGEN) or esiRNA (Sigma-Aldrich) using *Trans*IT-siQUEST reagent (Mirus Bio) following manufacturer’s protocol for 48-72h. Post transfection, these cells were used for qPCR, ATPlite assays and colony forming assays.

### Confocal Microscopy

OSC19-VC and TMEM16A cells were plated on coverslips and allowed to attach overnight. After formaldehyde fixation, cells were permeabilized in 0.1% triton (in PBS), blocked for one hour at room temperature and incubated for overnight at 4 degrees in primary antibodies against Lamp1 (Santacruz Cat#sc-20011) and Tomm20 (Invitrogen Cat# PA552843) at 1:1000 dilution. Next day, the cells were washed three times in PBS+0.05% tween and incubated in secondary mouse and rabbit antibody (Invitrogen) for one hour at room temperature. The secondary antibodies used were alexa flour 568 (red; Invitrogen #A11019) and 488 (green; Invitrogen A27034) at 1:5000 dilution. The cells are washed atleast 3 times in PBS+0.05% tween and stained with DAPI (Invitrogen #62248) for 10 minutes. The cells are washed and mounted in fluoro-gel (Electron Microscopy Sciences #17985-10).

Confocal z-stacks were collected using an Olympus Fluoview 1000 confocal microscope equipped with a 60x (1.4NA) optic, and analyzed using NIS-Elements (Nikon Inc., Melville NY). Co-localization was assessed by segmenting the lysosomal and mitochondrial signals based on intensity, size and shape. The percentage of total lysosomes containing mitochondria content was calculated using a binary “and” statement and expressed relative to total lysosomal count. At least 15 non-overlapping images were taken per experiment.

### Cell Proliferation Assay

Cells were plated in triplicate at a confluency of 2000-4000 cells/well in a 96-well plate. Following 24 hours, cells are treated as indicated. After 72 hours of treatment, cell proliferation was measured using WST-1 Reagent (Takara Bio, Inc., Clontech Laboratories, Inc.) as per manufacturer’s instructions. The absorbance was taken at 450 nm using the Epoch Plate Reader (Agilent BioTek).

### Colony Formation Assay

OSC19 were plated (in triplicate) in a 12-well plate at the confluency of 1000-2000 cells/well. After 24 hours, cells were treated at concentrations as described, and colonies were allowed to form until the control group reached near maximum confluency (at 7-10 days). Following treatment, cells were fixed in formalin (1:10 dilution) for 20 minutes and stained with 0.5% crystal violet for 20 minutes. Excess crystal violet was removed with tap water, and the plate was allowed to dry. The plate was then scanned and analyzed using the macro ‘colony area measurer’ of Image J.

### Western Blot

Cells grown in 10 cm dishes were collected after indicated treatments. Cell pellets were first lysed using RIPA buffer followed by sonification. For western blotting in HPV-tumors, ∼30mg tumor samples were minced on dry ice in cold RIPA buffer and homogenized using a pellet pestle motor followed by sonication. Samples were kept on ice for 15 minutes followed by centrifuging at maximum speed for 20 minutes. Protein quantification was performed from lysate (supernatant) using the Bradford assay, and 80 µg protein was loaded into gel. Actin normalization was done for each gel. Immunoreactive bands were visualized and quantified using the LiCor Odyssey system.

### Quantitative Real time PCR

Tissue samples from patients with SCCHN were collected after obtaining informed consent. All patients were subjected to biopsy before the initiation of any treatment. The clinical characteristics of all patient samples used for the study is listed in Table 2B. The tumor samples were transferred in RPMI media supplemented with antibiotics and stored in RNAse later until RNA isolation using the RNeasy Mini Kit (QIAGEN). Briefly, ∼25 mg tissue was disrupted in cold RLT buffer and homogenized through QIA shredder spin column. The lysate was again centrifuged before washing and eluting in water. RNA was quantified using the Gen5 Take 3 Module (Agilent BioTek) and assessed for quality with 260/230 absorbance ratio. For cell lines, cells were scraped into a pellet and RNA was isolated following Qiagen protocol. iScript Reverse Transcription (Bio-Rad) was used for cDNA synthesis. Quantitative PCR (qPCR) was conducted using SYBR green master mix using *ACTIN and GAPDH* as housekeeping controls. Relative gene expression was calculated using the 2^-ΔΔCq^ method. The primer sequences are listed in Table 3. PCR conditions used are 40 cycles at 15 sec for denaturation at 95°C, 30 sec annealing at 60°C, 30 sec extension at 72°C.

**Table 2A:**
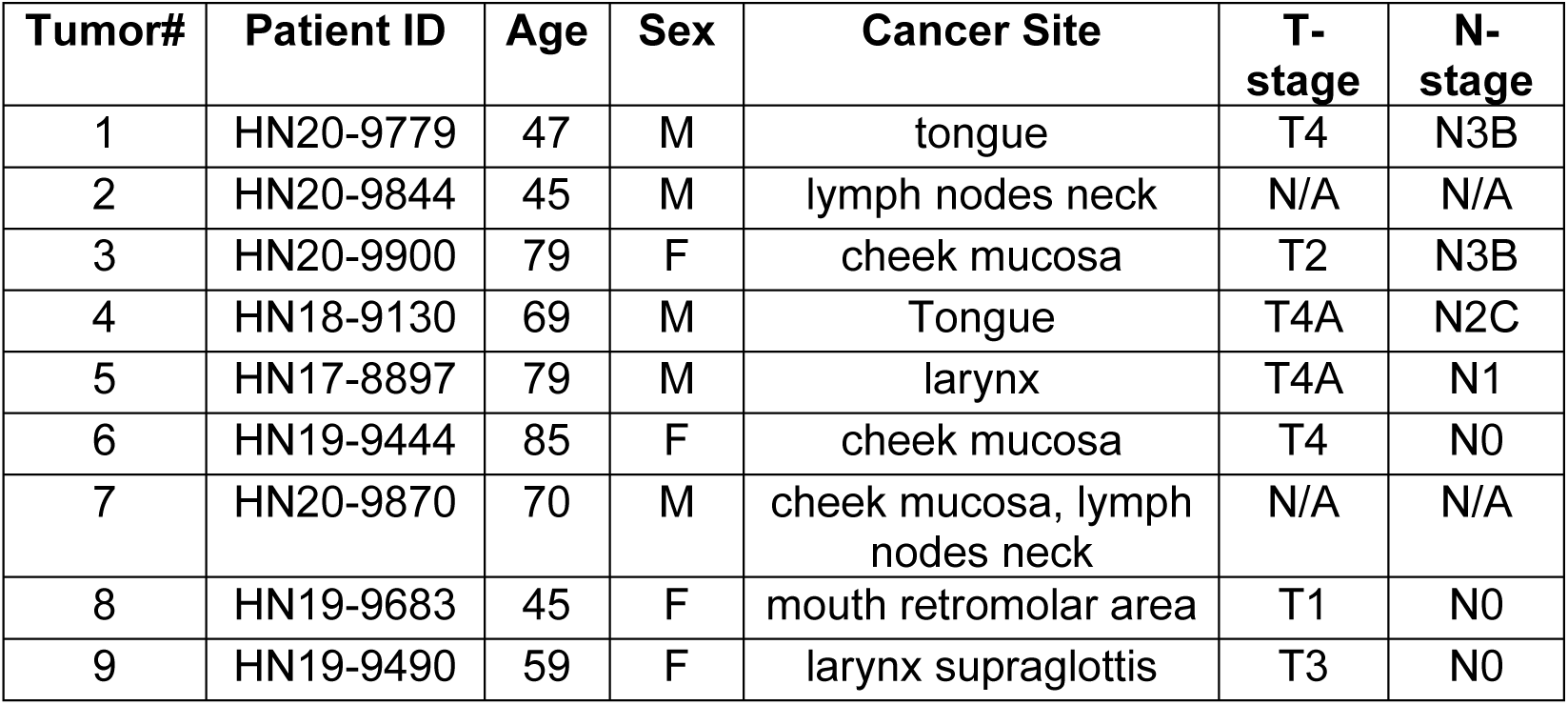
Clinical characteristics of patient samples used for western blotting.

**Table 2B:** Clinical characteristics of patient samples used for qPCR.

| **Tumor#** | **Patient ID** | **Age** | **Sex** | **Cancer Site** | **T-stage** | **N-stage** |
| --- | --- | --- | --- | --- | --- | --- |
| 1 | HN22-10407 | 65 | M | larynx | T3 | N0 |
| 2 | HN22-10390 | 61 | M | tongue | T2 | N0 |
| 3 | HN22-10389 | 74 | M | tongue | T3 | N0 |
| 4 | HN20-9844 | 43 | M | lymph node | T2 | N2C |
| 5 | HN19-9677 | 73 | F | mouth-overlapping lesion | T4 | N0 |
| 6 | HN20-9779 | 47 | M | tongue | T3 | N3B |
| 7 | HN22-10387 | 61 | M | hypopharynx | T4A | N2B |
| 8 | HN20-9742 | 78 | F | Maxilla | T4A | N3B |
| 9 | HN20-9753 | 64 | M | Mandible | T4A | N2A |
| 10 | HN20-9869 | 58 | F | Tongue | T4 | N2C |
| 11 | HN21-10045 | 74 | M | Tongue | T3 | N2B |
| 12 | HN21-10075 | 53 | M | Oropharynx | T4A | N3B |
| 13 | HN21-10099 | 61 | M | Tongue | T4 | N3B |
| 14 | HN22-10326 | 55 | F | Tongue | T3 | N2B |
| 15 | HN22-10397 | 47 | M | Larynx | T3 | N1 |
| 16 | HN22-10433 | 56 | M | Larynx | T4 | N1 |
| 17 | HN22-10407 | 86 | M | Larynx | T3 | N0 |
| 18 | HN21-10031 | 70 | F | Mandible | T4A | N0 |
| 19 | HN21-10101 | 70 | F | Maxilla | T4B | N0 |
| 20 | HN21-10154 | 63 | M | Floor of the mouth | T4 | N0 |
| 21 | HN22-10373 | 55 | M | Tongue | T3 | N0 |
| 22 | HN21-10201 | 71 | M | Floor of the mouth | T4A | N0 |
| 23 | HN21-10094 | 62 | M | Larynx | T4A | N0 |
| 24 | HN22-10448 | 65 | F | Mandible | T4A | N0 |

**Table 3:**
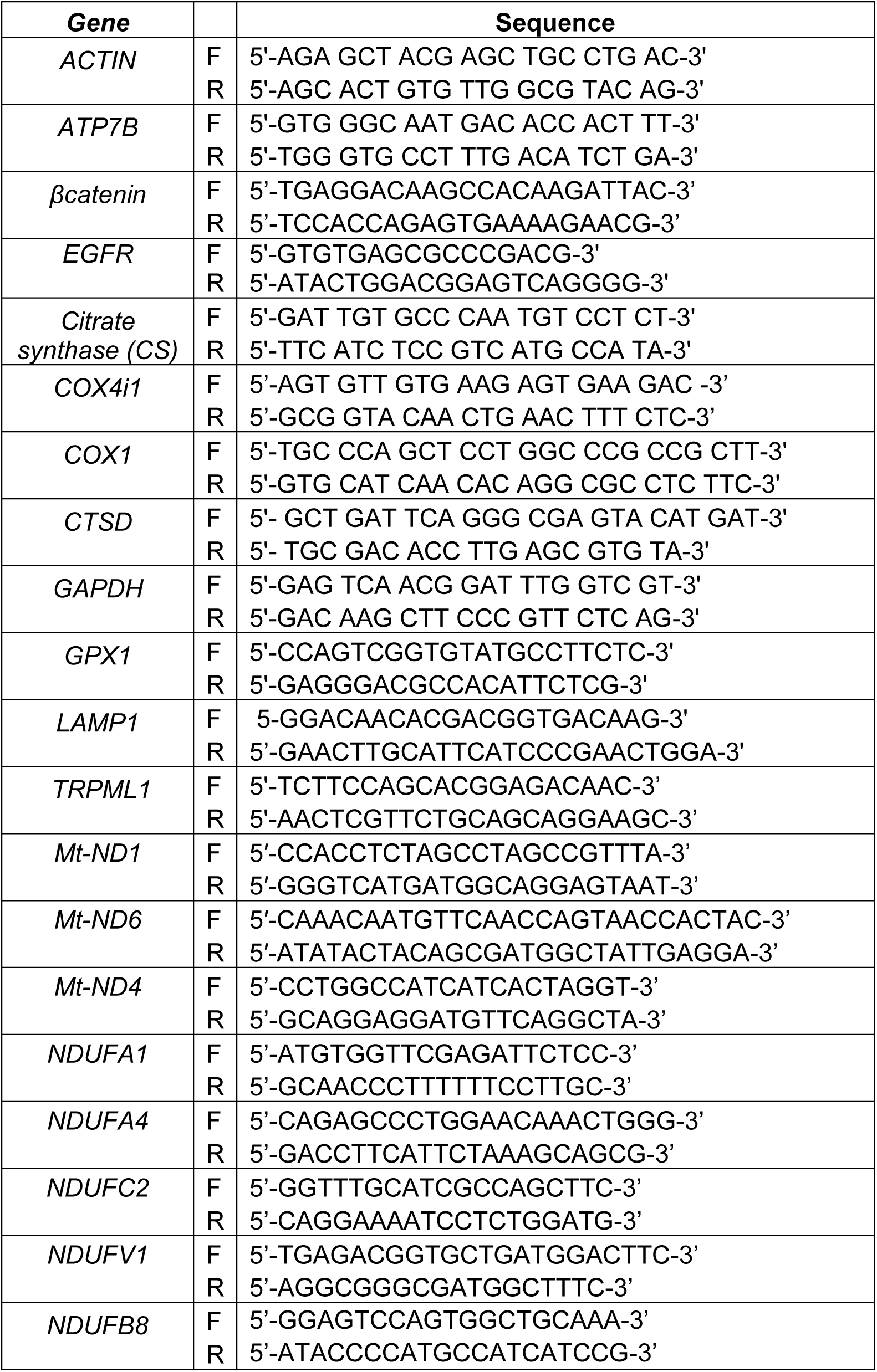
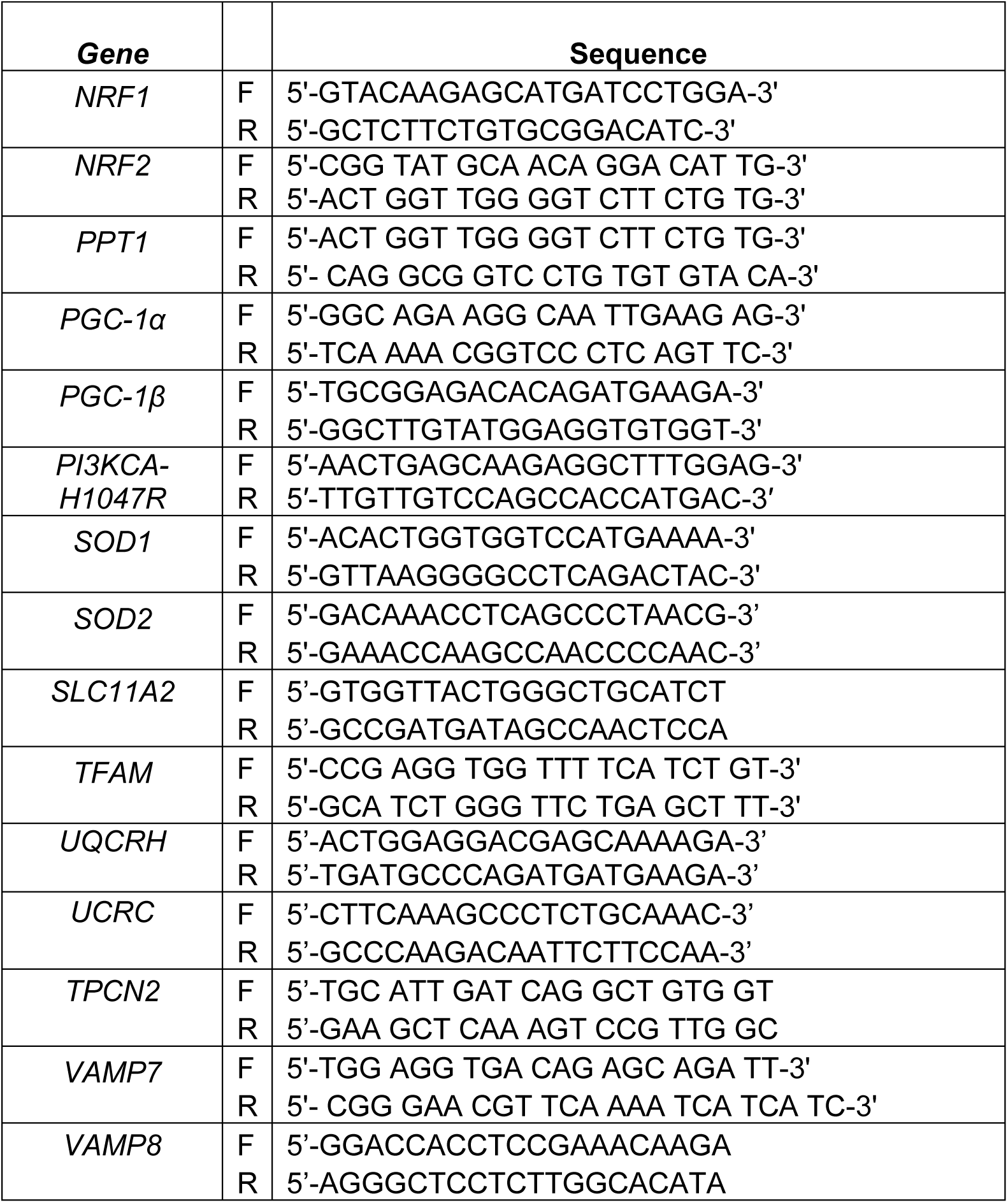
qPCR primers used in the study

### Mitotracker Green

Cells are plated in triplicate in 6-well plate and allowed to reach 70% confluency. After trypsinizing and washing once with PBS, ∼3e5 cells are suspended in 100nM mitotracker green (Invitrogen #M7515) diluted in cold Krebs buffer. A tube containing FCCP (5 µM) + mitotracker green is used as technical control for the experiment. Cells are allowed to stain on ice for 30 minutes. Tubes are centrifuged, washed once and resuspended in 250µl cold Krebs buffer and analyzed by flow cytometery on Beckman Coulter Cytoflex analyzer on 488nm laser. Non-viable cells were excluded from analysis using PI stain. Median fluorescence intensity (MFI) values were taken for unstained, mitotracker green stained and FCCP + mitotracker green stained groups.

### TOP/FOP assay

OSC19-VC and -TMEM16A were co-transfected with TOPflash or FOPflash plasmid along with the pLZ313-Renilla luc. The transfection was carried out with TransIT-X2 reagent (Mirus Bio.) in 6-wells. Post 48-hour transfection, 8,000 cells were plated/well in 96-well plate and Dual Glo Luciferase system (Promega) was used with a luminometer to measure expression levels in light units.

### Complex I Assay

Cell pellets were frozen on dry ice after collection and immediately submitted to University of Pittsburgh Center for Metabolism and Mitochondrial Medicine (C3M) for measurement of complex activity using spectrophotometric kinetic assay system. Briefly, complex I activity was measured by spectrophotometrically (340 nm) monitoring the oxidation of 100 μM NADH in the presence of 10 μM coenzyme Q_1_ in the presence and absence of 25 μM rotenone as previously described ^38^.

### ATP Production

At least 10,000 cells/well were plated in 96-well plate, allowed to attach overnight, and treated with indicated treatments. ATPlite Luminescence Assay System (Perkin Elmer Inc.) kit was used to measure ATP production. A standard curve of ATP was run and normalized using the crystal violet. Luminescence was read using the Synergy H1 Hybrid microplate reader (BioTek).

### Tetramethylrhodamine, ethyl ester (TMRE)

OSC19-VC/OE cells were plated at a confluency of 15000-20000 cells/well in a 96-well black plate. Following plating of cells, the quantification of changes in mitochondrial membrane potential was determined following the protocol of the TMRE-Mitochondrial Membrane Potential Assay Kit (Abcam). Briefly, live cells were first either stained with carbonyl cyanide 4-(trifluoromethoxy) phenylhydrazone (FCCP) or left unstained, then incubated with TMRE, and finally washed with 1X Phosphate Buffered Saline (PBS) and 0.2% Bovine Serum Albumin (BSA). The 96-black well plate was analyzed using the Epoch Microplate Spectrophotometer (BioTek) at excitation and emission values of 549 nm and 575 nm, respectively.

### *In vivo* Xenograft study

For MOC1 xenograft experiments, 5-week-old female C57BL/6J mice (Jackson Laboratory) were injected in bilateral flanks subcutaneously with 3e6 MOC1-VC and MOC1-TMEM16A cells (in 50% Matrigel). After tumors became palpable, mice were randomized into four treatment groups: MOC1-VC Control, MOC1-VC IACS, MOC1-TMEM16A Control, MOC1-TMEM16A IACS. Each group had 9-10 mice. IACS dissolved in 0.5% sodium carboxymethyl cellulose (Na-CMC) was administered to mice once a day via oral gavage for eight consecutive days (8 total treatments) at a dose of 10 mg/kg. Mouse weight (grams) and tumor volume (mm^3^) were taken daily. Tumor volume (mm^3^) was calculated using the equation: length(mm) x width(mm) x width(mm) / 2.

### In vivo PDX study

After surgical resection, the tumor sample was transported in RPMI media supplemented with Pen/Strep and anti-anti. The tumor was implanted into the flank of 6-week-old female NOD SCID mice and passaged once before being used for the study. The mice were randomized when average tumor volume reached an average of 100-150mm^3^. Mice were randomized into 2 groups of untreated and IACS treatment. The differences in relative tumor volume for each day was calculated using two-way ANOVA with Sidak’s multiple comparison test.

### Electron Microscopy of Xenograft tumors

Female nude mice were bilaterally injected with 3×10^6^ OSC19-VC and OSC19-TMEM16A cells until tumors reached 75-100 mm3. After sacrificing the mice, tumors were surgically removed and fixed in Karnovsky’s fixing buffer. The tumors were then sent to the Center for Biological Imaging at the University of Pittsburgh for further processing. Images were captured at 40000X. Image J software was used to analyze the mitochondria perimeter and area of the images.

### Statistics

All experiments are done atleast 3 times in triplicate, and combined data of N=3 is shown throughout the manuscript figures, unless otherwise stated. All data is represented as Mean ± S.E.M. Significance is defined as follows: *p<0.05, **p<0.001 and ***p<0.0001.

### Study Approval

HPV-tumors were obtained from the University of Pittsburgh Medical Center in accordance with established University of Pittsburgh IRB guidelines. A written informed consent was obtained from all the patients before inclusion in the study. The clinical characteristics of all patient samples used for the study is listed in Table 2. All animal studies were done in accordance with approved IACUC protocol.

## Acknowledgments

We thank Centre for Biological Imaging for help with quantifying confocal images. We are grateful to Dr. Satdarshan P. S. Monga for kind gift of TOP/FOP plasmids. We thank Mike Calderon for help with designing algorithm to analyze confocal images.

