## Supplemental Data for "Lysosomal mitochondrial interaction promotes tumor growth in squamous cell carcinoma of the head and neck"

Supplementary Figures and Legends

#### Supplemental Figure 1 : Lysosome-mitochondria interaction in TMEM16A cells is independent of mitophagy

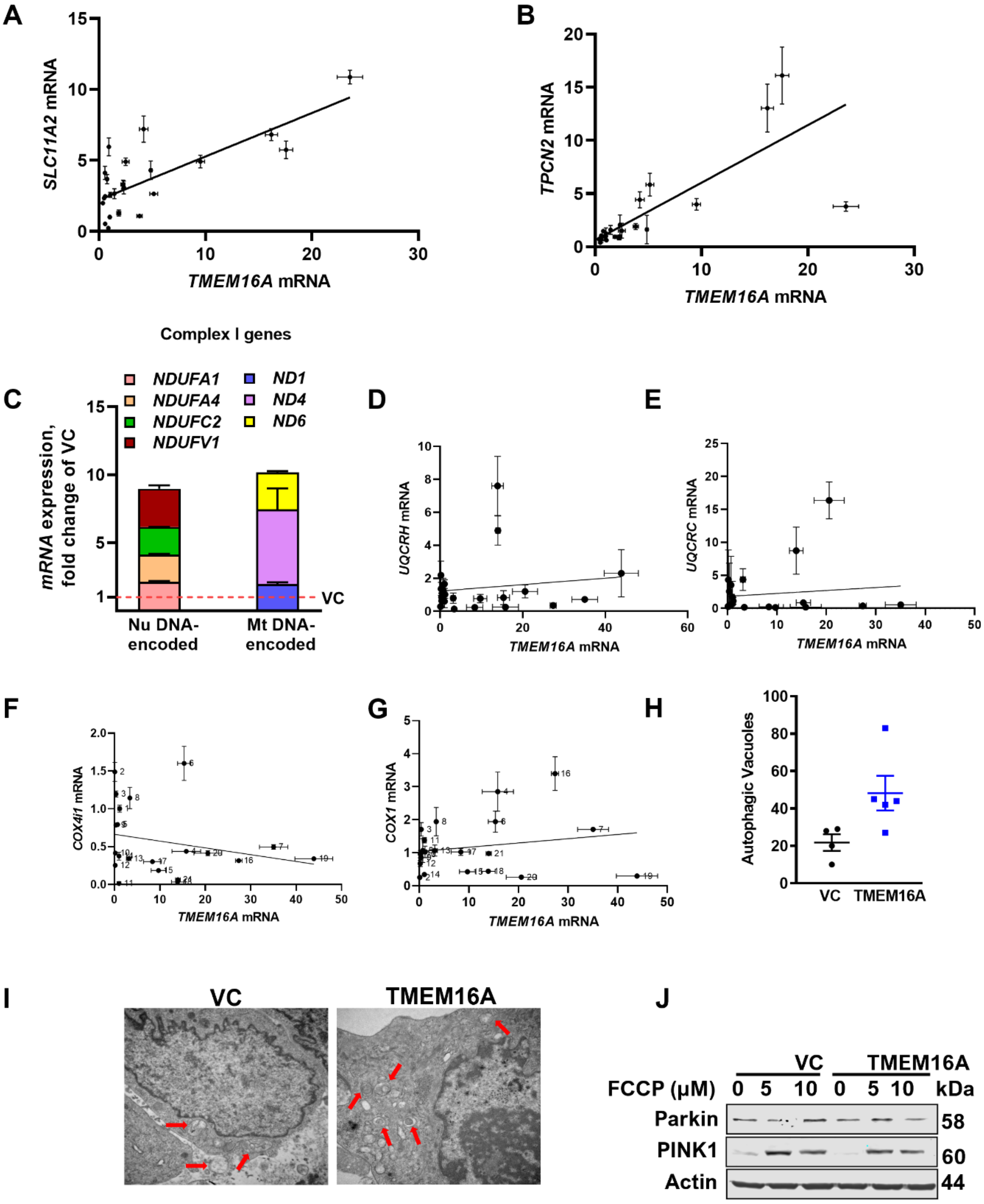

**Supplementary Figure 1: Lysosome-mitochondria interaction in TMEM16A cells is independent of mitophagy**

Correlation plots between TMEM16A expression and lysosomal gene expression in human SCCHN samples is shown in (A, B). The effect of TMEM16A expression on the expression of mitochondrial genes is shown in (C). Lack of correlation between TMEM16A and complex III gene expression (D,E) and with complex IV gene expression (F,G). The effect of TMEM16A on autophagy and mitophagy is shown in (H-J). Autophagy was assessed by measuring autophagic vacuoles. Mitophagy was evaluated by immunoblotting of PINK1 and PARKIN expression upon treatment with the mitochondrial decoupler, FCCP.

**Supplemental Figure 2: Wnt/β-catenin signaling has a role in lysosomal and mitochondrial biogenesis**

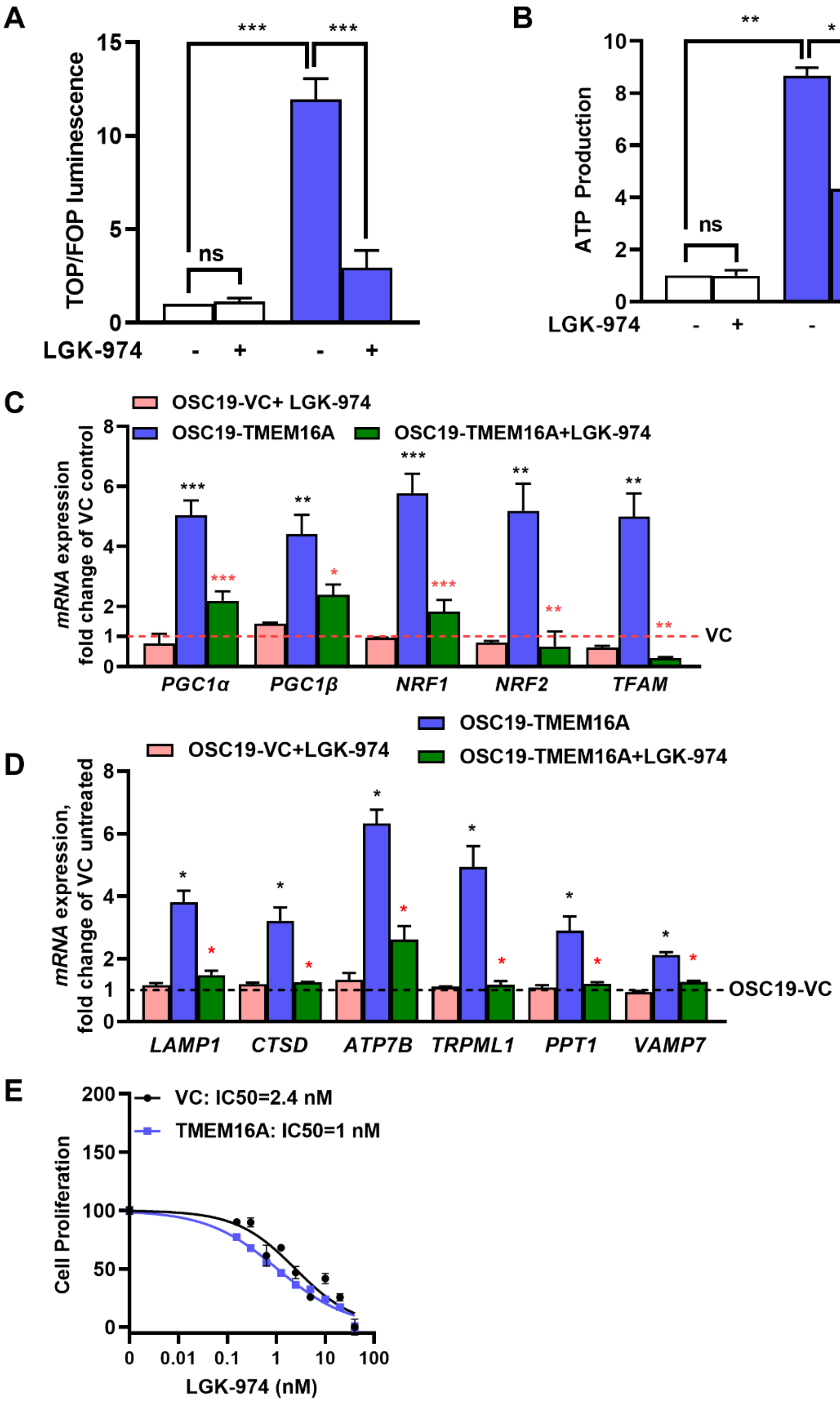

**Supplementary Figure 2: Wnt/β-catenin signaling has a role in lysosomal and mitochondrial biogenesis**

TMEM16A overexpression promotes β-catenin activity, that is abrogated by treatment with the porcupine inhibitor LGK-974 (1nM, 24h) (A). The effect of LGK-974 treatment on ATP production was measured using the ATPlite assay (B). The effect of LGK-974 treatment on markers of mitochondrial and lysosomal biogenesis is shown in (C ,D). The effect of LGK-974 treatment on cell proliferation, as determined by WST assay is shown in (E).

**Supplemental Figure 3: ETC uncoupler/ Complex III /IV inhibitors do not regulate lysosomal biogenesis**

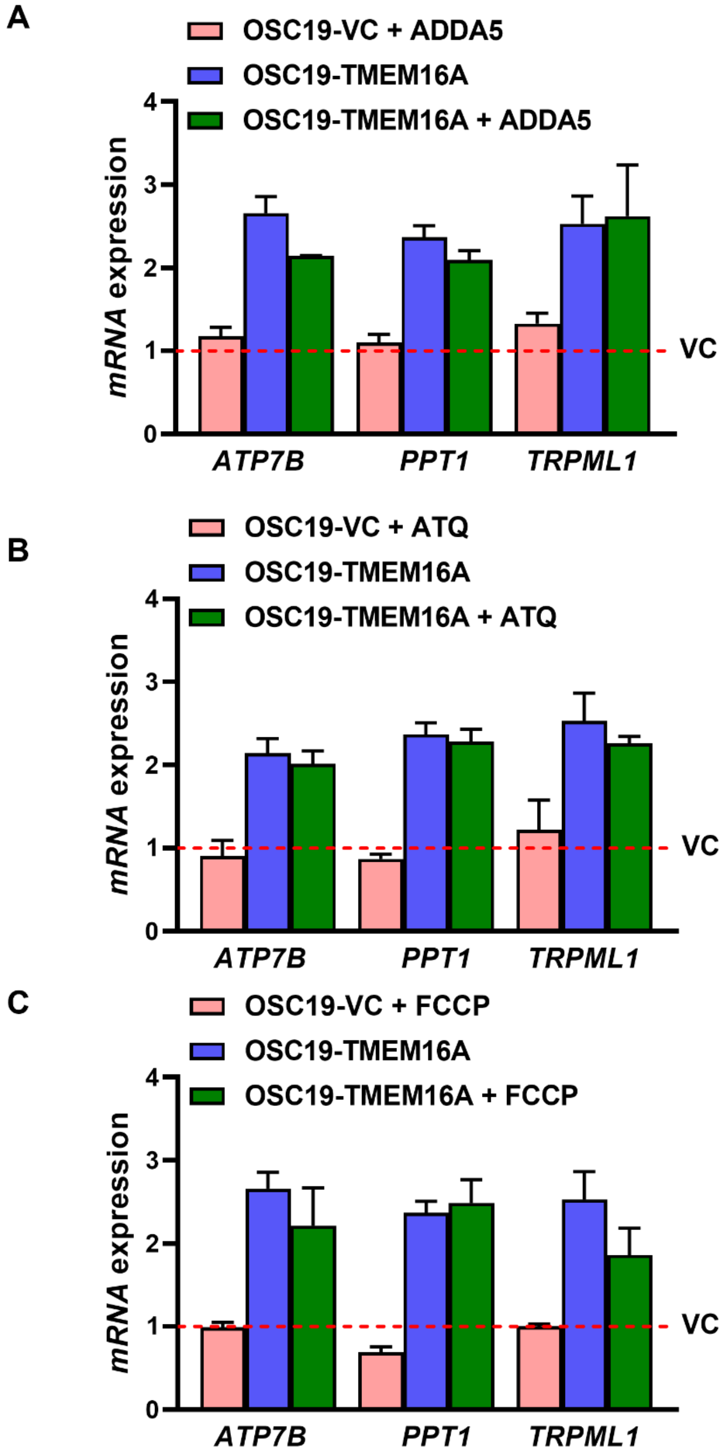

**Supplementary Figure 3: ETC uncoupler/ Complex III /IV inhibitors do not regulate lysosomal biogenesis**

Inhibition of mitochondrial complex III or IV does not impact lysosomal biogenesis. OSC19 cells were treated with ADDA5 (complex IV inhibitor) and assessed for lysosomal biogenesis (A). These cells were treated with Atovoquone (Complex III inhibitor) and assayed for lysosomal biogenesis (B). The effect of treatment with a mitochondrial decoupler, FCCP, is shown in (C).

**Supplemental Figure 4: Lysosomal mitochondria interface (LMI) is specific to complex I**

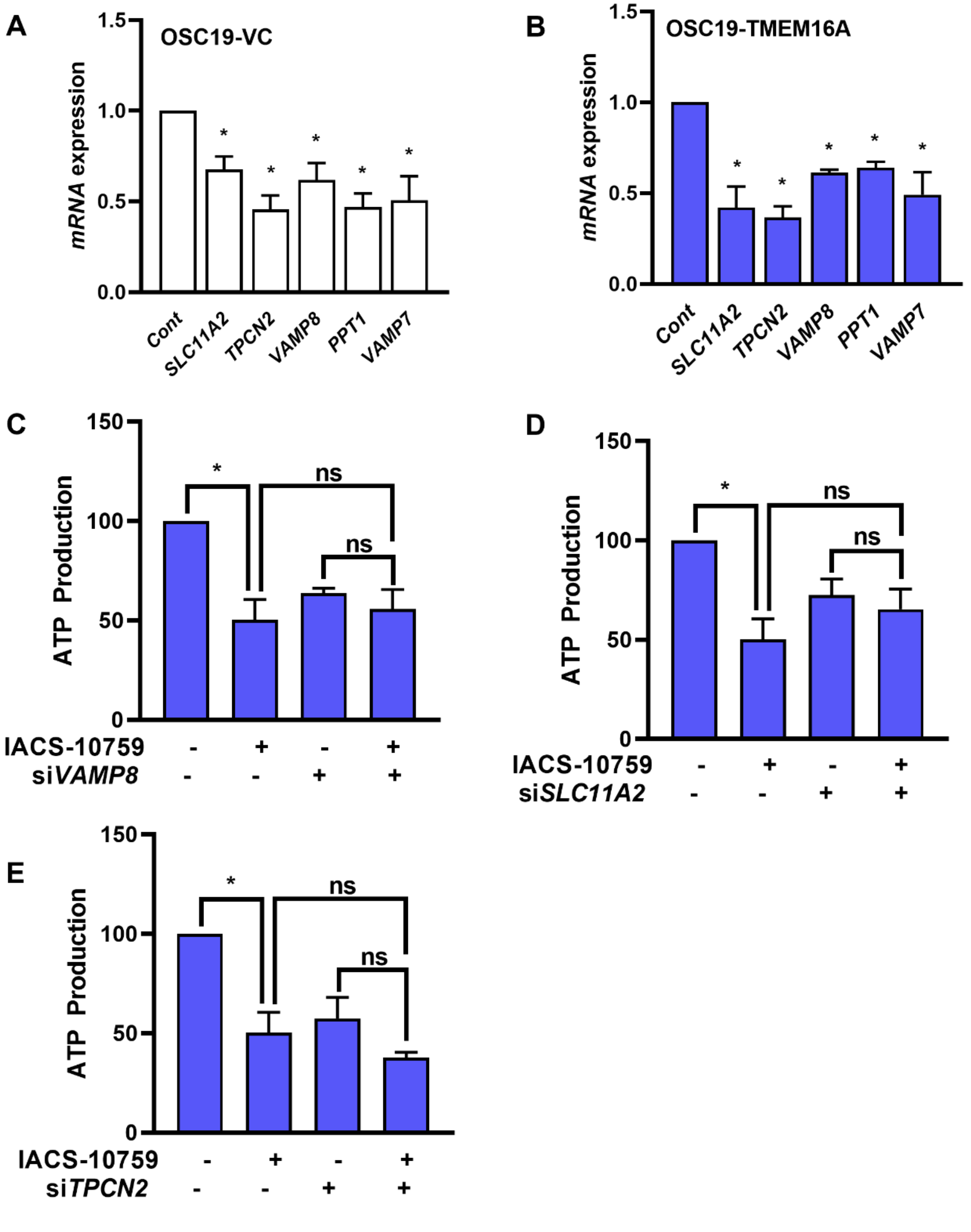

**Supplementary Figure 4: Lysosomal mitochondria cross talk (LMI) is specific to complex I**

The effect of siRNA treatment on gene expression of the enumerated genes in OSC19 VC and TMEM16A cells (A, B). Addition of IACS-10759 to siRNA targeting VAMP8, SLC11A2 and TPCN2 has no additional depletion of ATP production (C-E).

#### Supplemental Figure 5: OXPHOS inhibition is detrimental to *in vivo* growth in high-TMEM16A cells

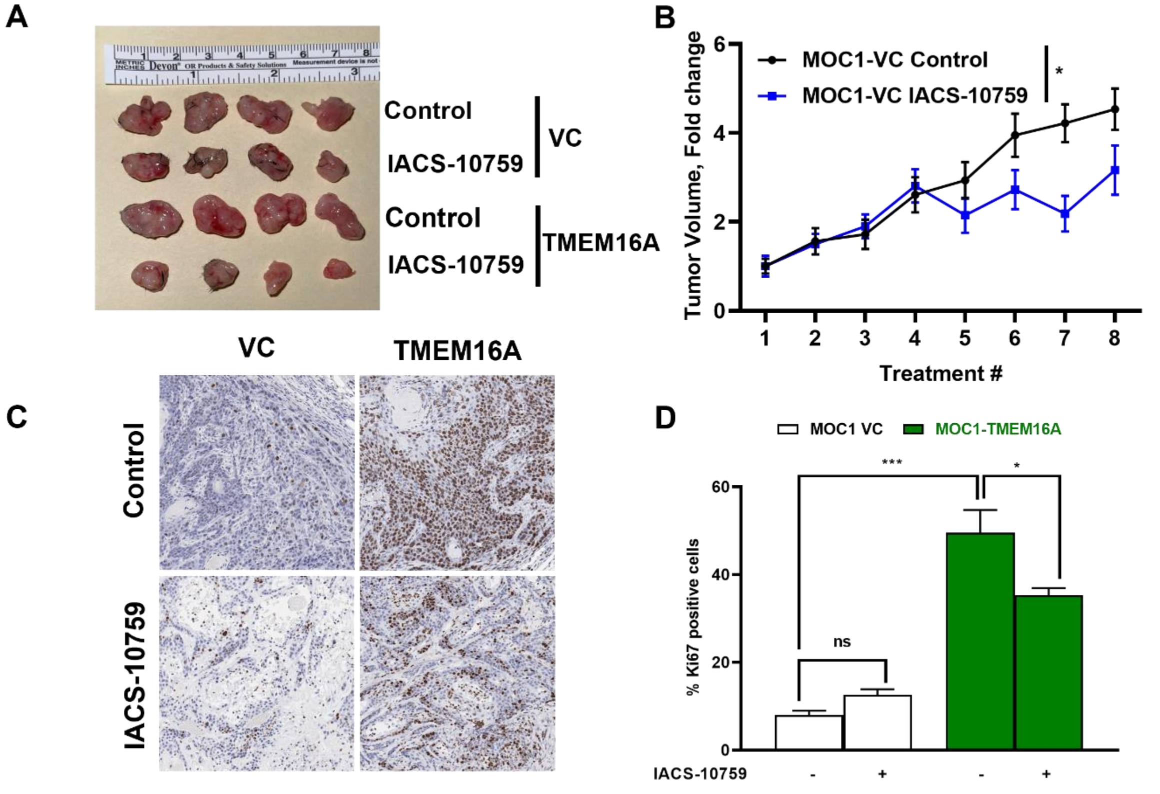


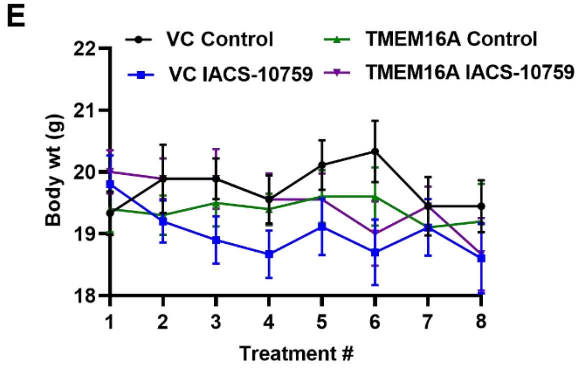


F

| Age, Sex | Cancer Site | T-stage | N-stage | Tx |
|---|---|---|---|---|
| 75y F | Cheek | T2 | N3B | Surgery |

**Supplemental Figure 5: OXPHOS inhibition is detrimental to *in vivo* growth in high-TMEM16A cells**

TMEM16A overexpression sensitizes tumor cells to complex I inhibition. MOC1 cells stably overexpressing TMEM16A were grown as subcutaneous xenografts and treated with placebo (control) or IACS-10759. Necropsy pictures are shown in (A). Growth curves for MOC1 control cells are shown in (B). Representative histology sections (Ki-67 staining) of tumors and quantitation are demonstrated in (C, D). The body weights of mice treated with IACS-10759 or diluent is shown in (E). Panel (F) presents the clinical characteristics of the patient from whom the PDX was generated.

### Supplemental Figure 6 : Mitochondrial function is diminished in TMEM16A knock-down model of HN30

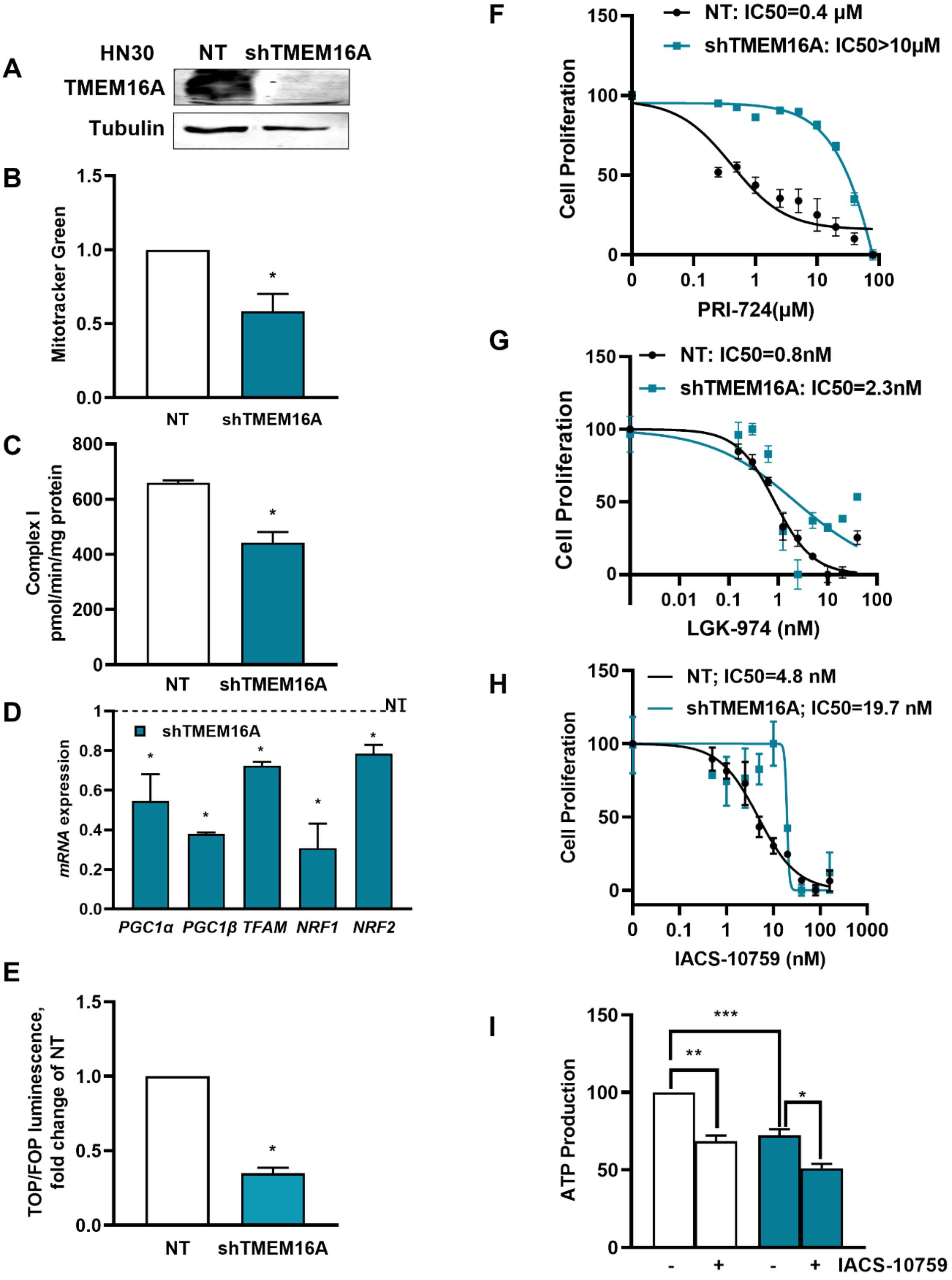

**Supplementary Figure 6: Mitochondrial function is diminished in TMEM16A knock-down model of HN30**

TMEM16A knock-down reduces mitochondrial respiration and de-sensitizes cells to IACS-10759. HN30 SCCHN cells were stably transfected with either non-targeting (NT) or shRNA to knock-down TMEM16A (shTMEM16A). Western blot indicating TMEM16A knockdown in the shTMEM16A cells (A). Mitochondrial mass was measured using MitoTracker Green (B). Complex I activity in these cells is shown in (C). Mitochondrial biogenesis is demonstrated in (D). TMEM16A knock-down resulted in reduced β-catenin activity (E) and reduced sensitivity to PRI-724 and LGK-974 induced cell death (F, G). TMEM16A knock-down also decreased sensitivity to IACS-10759 induced cell death (H). ATP levels are reduced upon TMEM16A knock-down, but the effect of IACS-10759 on reducing ATP production is diminished in cells devoid of TMEM16A (I).

**Supplemental Figure 7: Over expression of constitutively active PI3KCA or EGFR does not promote lysosomal or mitochondrial biogenesis**

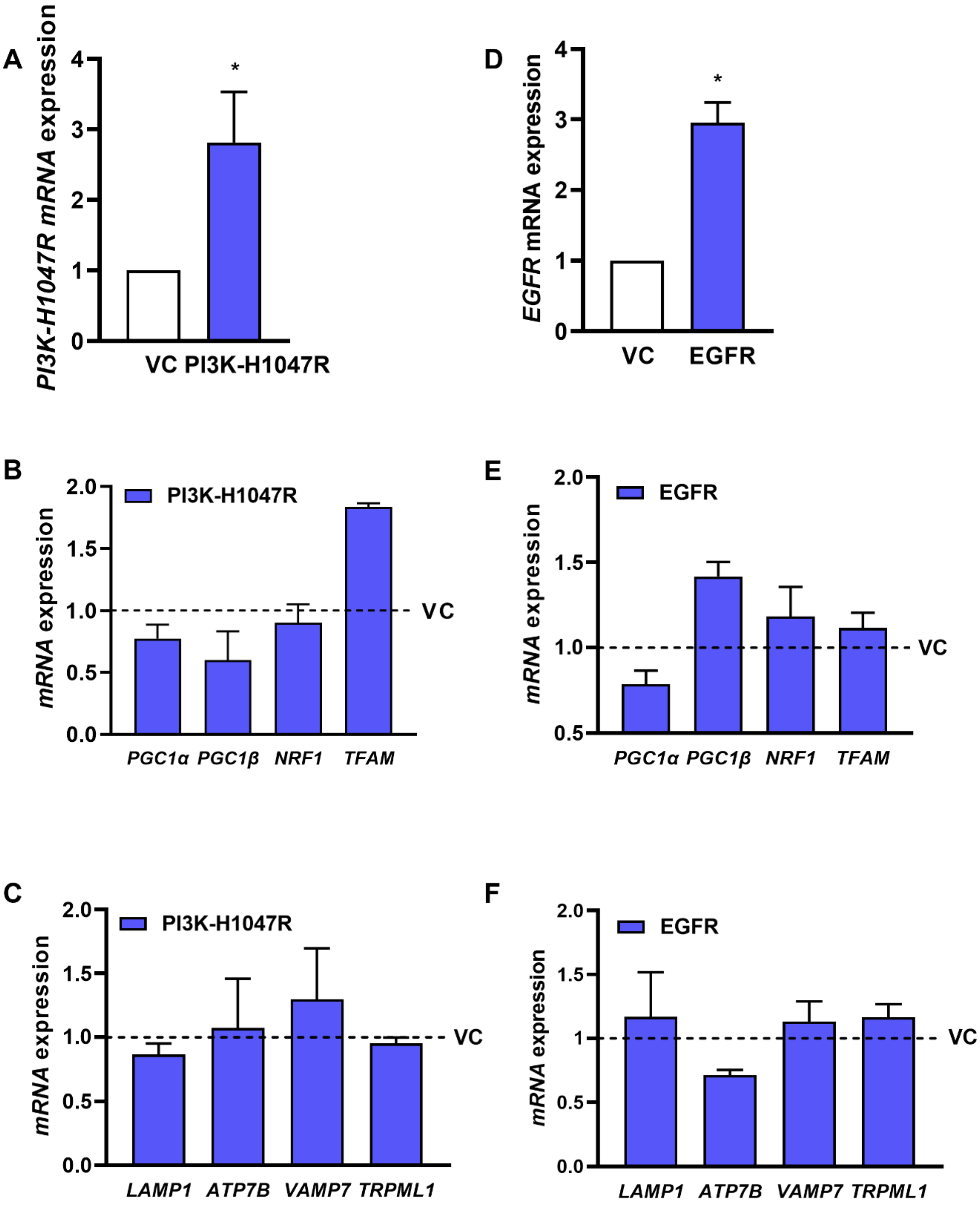

**Supplemental Figure 7: Over expression of constitutively active PI3KCA or EGFR does not promote lysosomal or mitochondrial biogenesis**

Gene expression analyses for mitochondrial and lysosomal biogenesis in OSC19 bearing transient overexpression of PI3K-H1047R (A-C) and EGFR (D-F).
